# Metabolic reprogramming by caloric restriction enhances acute phase virological control and reduces chronic inflammation in SIV-infected rhesus macaques

**DOI:** 10.64898/2026.03.11.711076

**Authors:** Naveen Suresh Babu, Chrysostomos Perdios, Maggie Hallmets, Anna T. Brown, Celeste Coleman, Christine M. Fennessey, Carolina Allers, Matilda J. Moström, Pratik Khare, Cissy Zhang, Brandon T. Smith, Nadia A. Golden, Adam Myers, Lara Doyle-Meyers, Alexandra Blaney, Robert V. Blair, Ahmad A. Saied, Ricki Colman, Brandon F. Keele, Anne Le, Clovis S. Palmer, Joseph C. Mudd

## Abstract

Nutrient metabolism influences HIV-1 replication, antiviral immunity, and chronic inflammation, yet is difficult to leverage for therapeutic gain. We sought to modulate metabolism in the non-human primate model of HIV-1 by caloric restriction (CR), a modality canonically known for its antiaging benefits. Four months of 30% CR was safe and resulted in broad and systemic metabolic reprogramming in healthy adult male and female rhesus macaques. Relative to that of *ad libitum-*fed animals, CR lowered the frequencies of target CCR5+ CD4 T cells in the gut mucosa. Upon infection with SIV, CR reduced acute phase viremia, dampened type I interferon signaling, and overall permitted a more vigorous cycling of CD8+ T cells in lymphoid tissues. CR-induced protection from SIV was associated with a robust up-regulation of glycolysis, which supported an early reduction in viremia that ultimately waned over time. During virologic suppression with antiretroviral therapy (ART), CR significantly limited gastrointestinal (GI) immune activation, improved tricarboxylic acid cycle flux, and lowered concentrations of soluble CD14 and several TNF-related molecules in plasma. Blood SIV DNA levels however were unchanged by CR, suggesting that residual GI dysfunction and inflammation can be decoupled from viral persistence. Our findings highlight that a dietary modality can limit pathology in a primate lentiviral infection. They also reveal the robust but temporally constrained nature of glycolysis in supporting an acute antiviral response.

**SIGNIFICANCE:** Caloric restriction (CR) is a safe dietary intervention known to confer anti-aging and health benefits across diverse animal models. However, its application in the context of infectious diseases has yielded mixed outcomes and has largely been limited to murine systems. In this study, we therefore employed CR to examine the impact of dietary modulation on SIV infection outcomes. Our findings demonstrate that CR reduced acute-phase viremia and attenuated markers of chronic inflammation following ART, effects that were associated with distinct metabolic signatures. Collectively, these findings underscore the importance of diet and nutrition in shaping chronic viral infection outcomes, such as SIV, within a clinically relevant non-human primate model.

## INTRODUCTION

Despite the profound success of combined antiretroviral therapy (ART) in durably suppressing HIV-1 replication, adverse residual pathologies persist in persons with HIV-1 (PWH). These include an incomplete repair of the gastrointestinal tract that is damaged during untreated HIV-1 infection^1–3^, which in long-term ART-suppressed subjects is linked to several inflammation-related comorbidities^4^. In addition, HIV-1-specific T cells are dysfunctional relative to T cells of other viral specificities and exhibit incomplete maturation and limited effector potential^5–7^. Lastly, the ability of HIV-1 to persist in latent form necessitates lifelong ART and the need to manage these complications long-term^8^.

Both the pathogenesis of HIV-1 and the host immune response are centrally regulated by cellular metabolism. Metabolic balance between processes such as glycolysis and mitochondrial oxidative phosphorylation (OXPHOS) are critical in supporting effective antiviral immunity^9–14^. These same metabolic processes, however, are also co-opted by replicating HIV-1 to sustain the viral lifecycle^15–18^, and chronic exposure to viral antigen in the absence of ART leads to the up-regulation of inhibitory immune checkpoint markers on virus-specific T cells and the loss of proliferative and effector potential. ‘Exhausted’ T cells that progressively dominate chronic HIV-1 display distinct metabolic profiles, characterized by a strict reliance on glycolysis at the expense of impaired mitochondrial function^19,20^. Importantly, immune metabolic abnormalities are not fully reversed despite virologic suppression with ART. For example, *in vivo* glucose uptake remains abnormally high in ART-suppressed PWH and associated with ongoing immune activation and residual inflammation ^21,22^. These data reinforce a generalized trend that chronic glycolytically dominated profiles are often associated with worse outcomes in ART-suppressed HIV-1, with more favorable outcomes associated with oxidative metabolism and improved mitochondrial health^23–26^.

Similar to other disease settings^23^, therapeutic HIV-1 treatment and cure strategies focused on specific metabolic pathways have faced both toxicity and efficacy hurdles^27,28^. Nutrition represents a pleiotropic and potentially safer approach to modulating the host metabolic and immunologic response to infections. The profound impact of nutrition on the host response to HIV-1 is highlighted by the fact that diets rich in saturated fats accelerate SIV disease progression in nonhuman primates, even in species that are non-pathogenic hosts of SIV^29,30^. While these studies are informative from a mechanistic standpoint, whether nutrition can be harnessed for therapeutic benefit in HIV-1 is largely unexplored. Caloric restriction (CR) represents one of the most robust nutritional interventions. Reducing caloric intake without malnourishment has been shown to lower the incidence of aging-associated pathologies across a diverse array of species^31,32^. In adult non-obese mice, the unique metabolic state imparted by CR can also confer enhanced protection against a range of pathogens^33–35^. Nonetheless, it is currently unknown whether tuning caloric intake can optimize host immunity in higher-order mammals.

In this study, we safely regulated daily calorie intake within a time-restricted feeding window in healthy male and female rhesus macaques. We then infected animals with SIV and then durably suppressed viremia with ART. We provide evidence that CR provoked a glycolytically skewed response to SIV that induced a robust but ultimately transient virologic protection. During ART, CR significantly alleviated gut mucosal immune dysfunction and was overall anti-inflammatory. Our data reveals a broad anti-inflammatory effect of CR in a primate lentiviral infection and uncovers a context-dependent nature of glycolysis to acute SIV protection.

## RESULTS

### CR is safe and induces modest weight loss in healthy non-obese rhesus macaques

To implement CR, we enrolled six healthy adult, non-obese, male and female Indian rhesus macaques (**Table 1**). The median age of the animals was 11.8, corresponding to a human equivalent of 30-40 years (**Table 1**)^36^. Chow, which consisted of standardized high-fiber/low-fat condensed ‘biscuits’, was provided *ad libitum* (AL) to RMs in singly housed conditions, and the number of biscuits consumed by each animal was recorded daily by the TNPRC veterinary staff (**Fig. 1A**, AL phase). In order to balance accurate AL reference appetite determination with the need to provide social enrichment, after 90 days, animals underwent a ‘hybrid-housing’ protocol in which food was made available to animals between the hours of 8:00-16:00 in singly housed conditions (**Fig. 1A**, time restriction phase). At 16:00 hours, biscuits were counted, food was removed from the cage, and animals were socially pair-housed without food for the remainder of the day. This routine was repeated daily and introduced a time-restricted (TR) feeding element to the study, which has been noted to optimize the health benefits of CR^37^. TR enforced a minimum fasting time of 16 hours (**Fig. 1B**) but induced variable changes to caloric intake that were not consistent in one direction across study animals (**Fig. 1C**). When compared to enrollment, 2 months of TR resulted in modest reductions in body weight (**Fig. S1A**). Levels of blood glucose, triglycerides, and cholesterol, measured as part of a routine blood chemistry panel, remained unchanged during TR (**Fig. S1B–C).** Similarly, immune phenotypes of circulating memory CD4⁺ and CD8⁺ T cells including markers of T cell activation were largely stable across time (**Fig. S1G-J)**, suggesting that the timed feeding and alteration to housing protocols had a limited effect on clinical metabolic and circulating immune cell phenotypes.

**Figure 1.**
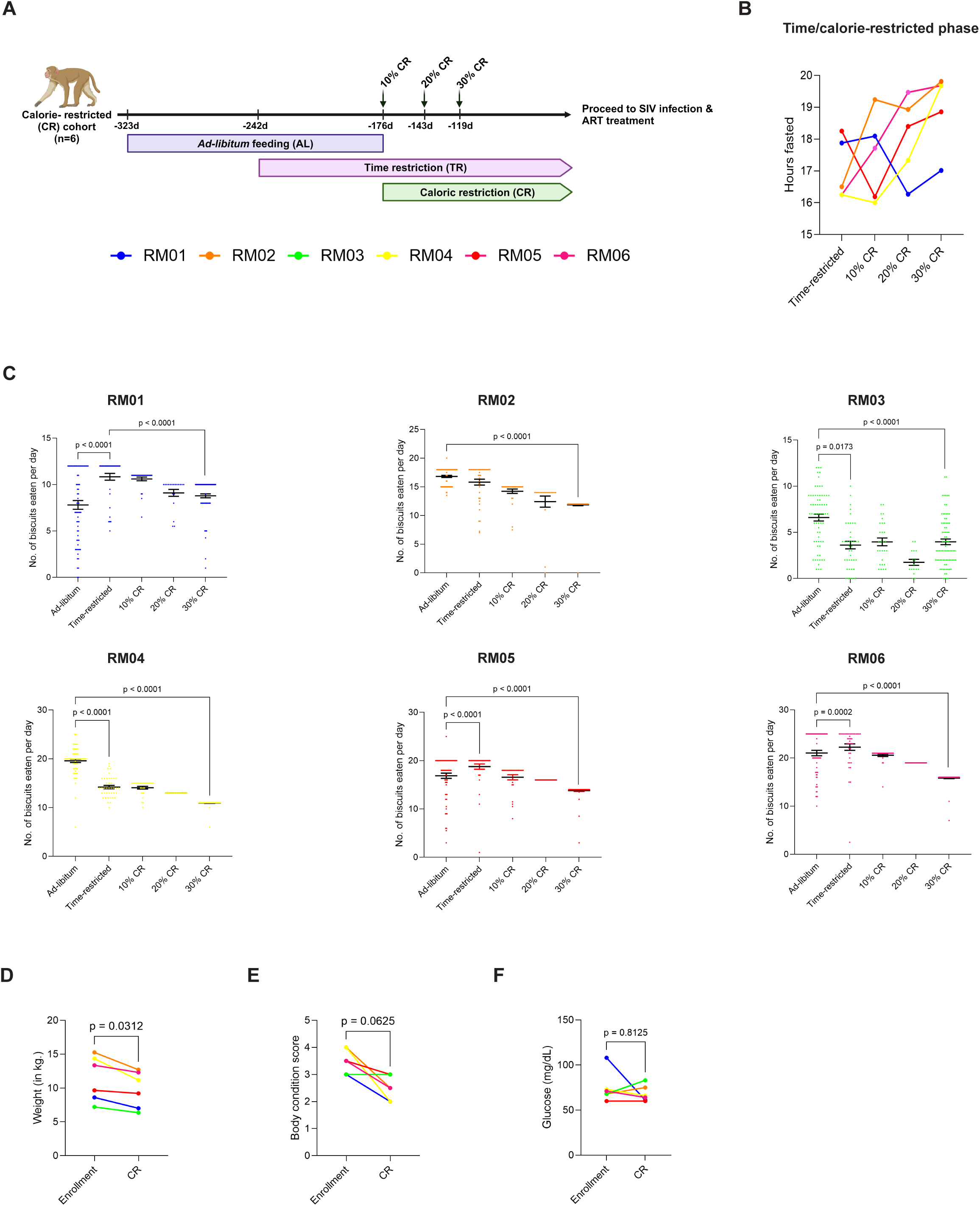
Implementation of time and caloric restricted feeding regimens in healthy non-obese male and female rhesus macaques. **(A)** Schematic representation of the *adlibitum* (AL), time restricted, CR titration and maintenance phases of the study in rhesus macaques (n=6). Animals were maintained on time and caloric restricted feeding regimens for 4 months prior to SIV infection. **(B)** Number of daily hours the animals were fasted through time restricted, CR titration and maintenance phases and caloric restriction. **(C)** Summary data of Number of biscuits consumed during each day of the respective feeding phase in individual animals. **(D)** Weight and **(E)** body condition scores at study enrollment and 3 months after maintaining 30% CR. **(F)** Blood glucose levels at study enrollment and 3 months after maintaining 30% CR. **Statistical analyses: (C-G)** Wilcoxon matched-pairs signed rank test. α=0.05.

**Table 1:**
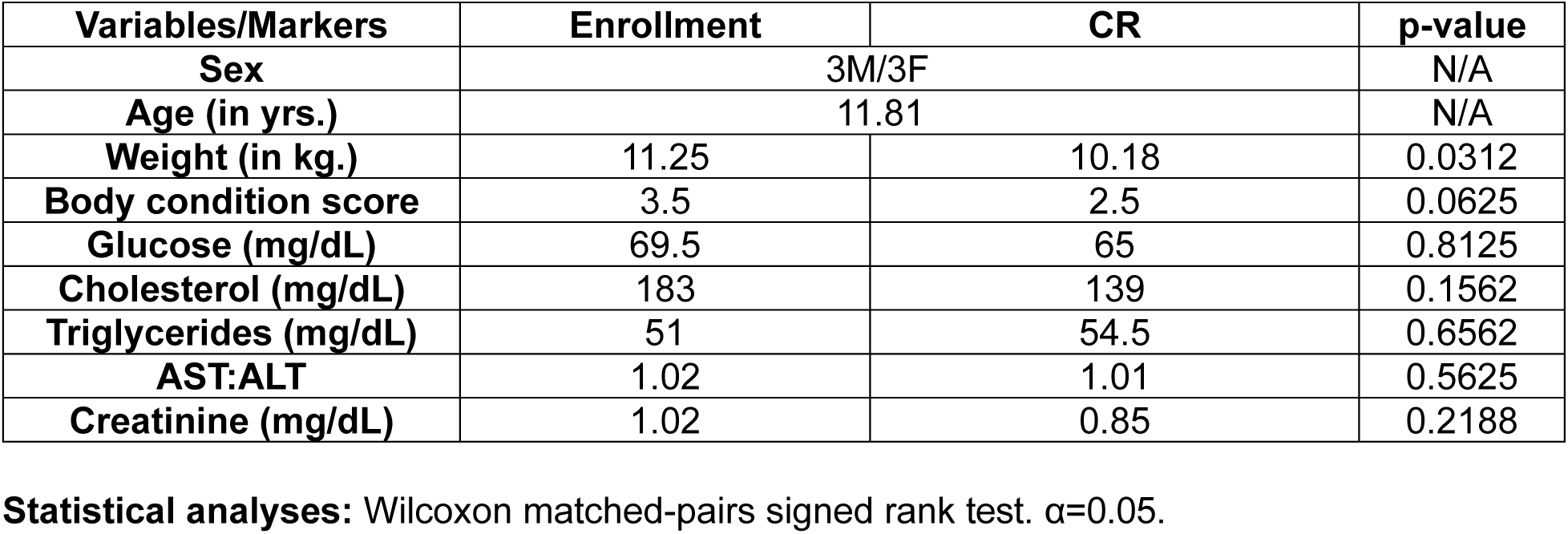
Median age, weights, BCS, and plasma concentrations of key physiological markers of the CR cohort (n=6) at enrollment and following 30% caloric restriction.

Following 2 months of TR, the number of daily biscuits supplied to the animals was restricted within the TR-feeding window by monthly 10% intervals until a target restriction of 30% was attained relative to each animal’s *ad libitum* reference (**Fig. 1A**, CR phase). The implementation of CR within the feeding window caused some animals to eat their food more rapidly (RM02, RM04, RM06) (**Fig. 1B**), consistent with a self-enforced fasting period noted in murine studies of CR^37,38^. One animal (RM03) naturally consumed the majority of its food in the evening and hence was not compliant with a daytime feeding schedule; this animal was thus caloric- but not time-restricted and remained singly housed through the remainder of the study.

Restricting daily biscuits by 30% predictably resulted in significant reductions in daily caloric intake relative to baseline food intake (**Fig. 1C**). 30% CR maintained for 4 months caused a modest but significant reduction in weight from enrollment levels (**Fig. 1D**) and also tended to reduce animal body condition scores (BCS) (**Fig. 1E**), a qualitative measure of fat/lean mass ratio (5 = morbidly obese, 1 = emaciated)^39^. Despite modest weight loss, cholesterol, triglycerides, and markers of liver and kidney function were not significantly altered by CR (**Table 1**). We also investigated the impact of CR on measures of blood sugar homeostasis. Despite the reduction of calorie intake, CR did not alter levels of fasting blood glucose relative to these levels at enrollment (**Fig. 1F**). Overall, in a healthy cohort of male and female rhesus macaques, 4 months of CR within a TR feeding window resulted in modest weight loss but did not systemically alter major clinical panel metabolites.

### CR enhances virologic control during acute SIV infection

To determine cross-sectionally how manipulating the timing and quantity of food intake would impact the immune and metabolic response to SIV, following 4 months of maintaining CR we introduced a separate control cohort of 11 rhesus macaques that received continuous, *ad libitum* (AL) access to the same high-fiber/low-fat condensed biscuits as CR animals (**Fig. 2A**). The age, sex, weight and BCS at enrollment among AL and CR groups were not significantly different (**Table 2**). We also profiled clinical serum metabolites and cellular immune phenotypic markers of activation, proliferation, and maturation of lymphoid and myeloid cells across multiple anatomic sites prior to SIV (gating strategy, **Fig. S2**). After 4 months of CR, animals exhibited BCS lower than the AL-fed control group; however, weights and clinical metabolite levels were comparable (**Table S1)**. We also considered the possibility that manipulating the timing and quantity of food intake would lead to increased physiological stress within the CR cohort given that cortisol, a stress-induced hormone, is known to be elevated by CR ^40–43^. We thus measured plasma concentrations of four corticosteroids (Cortisol, Cortisone, Corticosterone, and 11-Dehydrocorticosterone) by mass spectrometry in AL and CR animals just prior to SIV infection and found no group differences in these levels (**Fig. S3A-D**), suggesting a lack of stress-induced phenotypes in the CR cohort. Finally, we performed principal component analysis (PCA) of 15 immune phenotypic markers (see, gating strategy, **Fig. S2**) across blood, lymph node, and colon to determine immunological differences between AL and CR cohorts prior to SIV infection. Overall, the global immunologic profiles across tissues did not differ between AL and CR groups (**Fig. S4A**). Notably, activation markers HLA-DR and CD69 memory CD4^+^ and CD8^+^ T cells did not differ in blood, gut, or lymph nodes prior to SIV infection (**Fig. S4B-G**).

**Figure 2.**
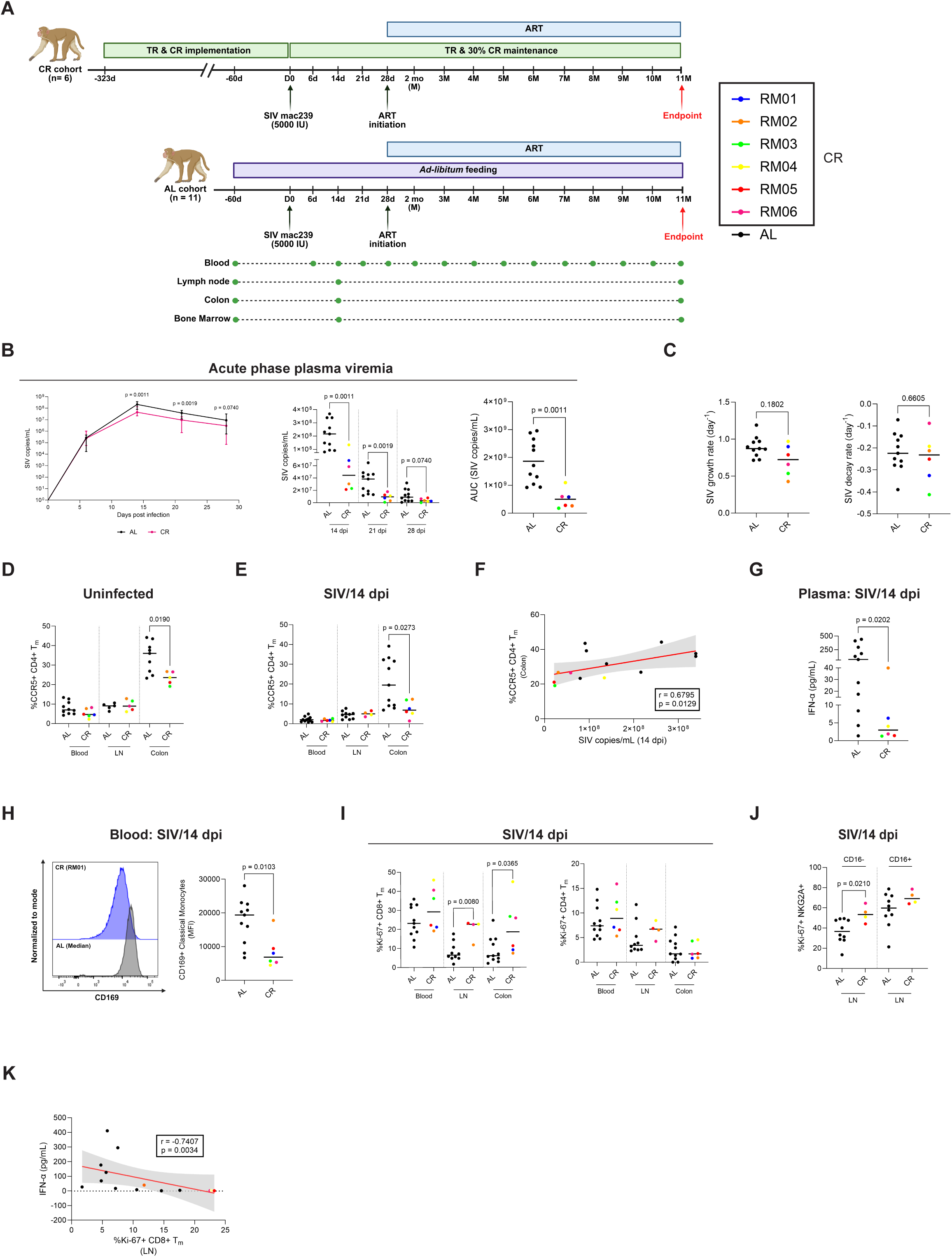
Preventative CR enhances virologic control of peak viremia. **(A)** Schematic depicting the timeline of SIV infection and ART initiation in AL (n=11) and CR animals (n=6). **(B)** Plasma viral load (log10 SIV copies/mL) in AL and CR animals at longitudinal timepoints of 7-, 14-, 21-, and 28-days post-infection (dpi) along with area under the curve analysis of viremia across the first 28 days of SIV. VL AUC was calculated using the trapezoidal method. **(C)** Dot plots depicting SIV growth (0-14 dpi) and decay (14-28 dpi) rates per day calculated from plasma viral load. **(D & E)** Percentage of CCR5^+^ CD4^+^ and CD8^+^ total memory T cells in AL and CR animals across blood, pLN and colon at uninfected **(D)** and 14 dpi **(E)** time points. **(F)** Two-sided spearman correlation between %CCR5^+^ CD4^+^ memory T cells in the colon and plasma viral load (SIV copies/mL) at 14 dpi. **(G)** Quantification of plasma IFN-α (pg/mL) in AL and CR animals at 14 dpi through ELISA. **(H)** Mean fluorescence intensity (MFI) of CD169 expression in CD14+ CD16- classical monocytes in the blood of AL and CR animals at 14 dpi (right) with a representative histogram from one animal in each cohort (left). **(I)** Frequencies of Ki-67^+^ CD4^+^ and CD8^+^ total memory T cells in AL and CR animals across blood, pLN and colon at 14 dpi. **(J)** Frequencies of Ki-67^+^ CD16^+^ and CD16^-^ NK cells in the pLN of AL and CR animals at 14 dpi through flow cytometry. **(K)** Plots depicting two-sided Spearman correlation between plasma IFN-α and frequencies of Ki-67^+^ CD4^+^ memory T cells in the pLN of AL and CR animals at 14 dpi. **Statistical analysis: (B-E, G-J)** Mann-Whitney U test and **(F & K)** Spearman non-parametric correlation with confidence bands generated through linear regression. α=0.05.

**Table 2:** Key demographics and levels of clinical metabolic analytes of AL and CR animals at study enrollment.

| **Parameters** | **Ad-libitum (AL)** | **Calorie-restricted (CR)** | **p-value** |
| --- | --- | --- | --- |
| **No. of animals** | 11 | 6 | N/A |
| **Sex** | 6M/5F | 3M/3F | >0.9999<sup>1</sup> |
| **Age** | 12.02 (5.80-18.43) | 11.81 (9.93-12.04) | 0.0983<sup>2</sup> |
| **Enrollment body condition score** | 3.75 (2.0-5.0) | 3.5 (3.0-4.0) | 0.9035<sup>2</sup> |
| **Enrollment weight (in kg.)** | 11.38 (6.55-17.0) | 11.25 (7.20-14.75) | 0.4936<sup>2</sup> |

**Statistical analyses:** 1: Fisher's exact test 2: Mann Whitney U-test. $\alpha=0.05$ .

We next infected both AL and CR cohorts with 5,000 infectious units of a barcoded SIVmac239M clone by single intravenous (i.v.) administration (**Fig. 2A**). Upon infection with SIV, CR transiently reduced acute phase viremia (**Fig. 2B**). The early period of exponential SIV replication between 0-7dpi was not different between groups, however CR reduced plasma viral load at 14- and 21-days post-infection (dpi) corresponding to peak viremia and the early emergence of adaptive immunity (**Fig. 2B**). Notably, the protective impact of CR was not sustained out to 28 dpi, as viremia tended to be lower in CR animals but was not statistically significant (**Fig. 2B**). Overall, CR resulted in ∼1 x 10^9^ fewer viral particles/ml across 28 days of infection as defined by area under curve (AUC) analysis (**Fig. 2B**).

To determine whether CR-induced virologic control reflected an imprint of heightened antiviral immunity, we first examined the viremia growth (0-14 dpi) and decay (14-28 dpi) rates between groups, reasoning that earlier and more robust immune responses would result in slower exponential growth and/or faster viremic decay. Relative to that of AL-fed animals, we found no evidence that CR altered SIV replication kinetics, including the early ramp-up growth rate (0-14 dpi, **Fig. 2C**) or the rate of plasma viremia decaying from peak to set point (14-28 dpi, **Fig. 2C**). To interrogate virologic dynamics in more granular detail, we longitudinally sequenced the unique barcode region of SIVmac239M virions in plasma across acute infection. SIV barcoding technology allows for the resolution of distinct SIV strains that comprise plasma viremia, and tracking their compositional changes across time can reveal signatures of immune selection pressure^44^. We assessed the community structure of distinct viral barcode strains by ecologic metrics, reasoning that heightened CR-induced immunity would result in a lower absolute number of unique viral barcodes replicating and more even proportional representation, preventing any single clonotype from achieving selective advantage and becoming overly dominant. Across 28 days of acute SIV we found no evidence that CR altered the number (richness) or the relative distribution of distinct replicating SIV strains (Shannon diversity, Gini clonality) (**Fig. S5A-C**). The data suggests that the protective effect of CR was due to a global ‘scaling down’ of viremia, and not due to pruning of select viral lineages by the immune system.

We next assessed the levels of target CCR5^+^ CD4^+^ T cells in blood and tissues, reasoning that more limited target cell availability would globally alter the size of the replicating pool, particularly in the gut that is target cell rich and a prominent anatomic site of early viral replication^2,45,46^. Prior to SIV, CR significantly reduced the frequencies of target cells in the colon (**Fig. 2D**), a finding unique to the gut and not observed at other anatomic sites of blood or lymphoid tissues (**Fig. 2D**). Moreover, colonic, but not blood or lymphoid memory CCR5^+^ CD4^+^ T cells remained significantly lower in CR animals at 14 dpi (**Fig. 2E**). Pre-infection frequencies of target cells in the colon correlated directly with the level of peak viremia at 14 dpi (**Fig. 2F**), suggesting that target cell restriction at gut mucosa sites was more strongly linked to CR-mediated reductions in acute phase VL.

Acute viremia is marked by a robust type-I interferon response that in progressive hosts is sustained across infection^47,48^. Thus, we next determined how CR influenced the type-I interferon response during acute SIV. Consistent with lower viremia, levels of IFN-α were reduced in the plasma of CR animals at 14 dpi (**Fig. 2G**). Surface expression of CD169, a canonical marker of type-I interferon exposure^49–51^, was also reduced on circulating CD14^+^ CD16^-^ classical monocytes (**Fig. 2H**). Although type-I interferon is essential in regulating protective antiviral programs, it is also anti-proliferative^52–54^, and persistently high IFN-α has been paradoxically linked to CD8⁺ T-cell exhaustion and reduced effector T cell proliferation^55,56^. To investigate the relationship between IFN-α exposure and the priming of antiviral immunity, we measured Ki-67 expression on memory CD8⁺ T cells as an indicator of cellular proliferation. At 14 dpi, CD8⁺, but not CD4^+^ memory T cells of CR animals exhibited higher Ki-67 expression in both the colon and peripheral lymph nodes (pLN) that are early anatomic sites of high viral burden (**Fig. 2I**). Increased cycling activity also extended to NKG2A^+^ CD16^-^ natural killer (NK) cells in the CR pLN (**Fig. 2I**). Importantly, we observed significant inverse correlations between plasma levels of IFN-α and the frequency of LN cycling CD8^+^ T cells (**Fig. 1K**). Taken together, the data indicates that lower SIV replication and a less inflammatory environment in CR animals was linked to less restrained proliferation of NK and CD8+ T cells in lymphoid tissues.

### CR enhances glycolysis during early acute SIV

To investigate whether CR-induced protection during acute SIV was linked to a distinct metabolic profile, we performed untargeted metabolomics in plasmas of CR animals and a subset of AL-fed animals (N=6) both prior to and 14 days post-infection with SIV. Out of a total of 208 detected metabolites, 19.7% and 18.2% of these met non-FDR corrected significance at baseline and 14 dpi, respectively (**Fig. 3A**). Metabolites that became differentially abundant (DA) during acute SIV in both AL and CR groups involved metabolites relating to the processing of amino acids, the most prominent of which being tryptophan metabolism (**Fig. 3B**).

**Figure 3.**
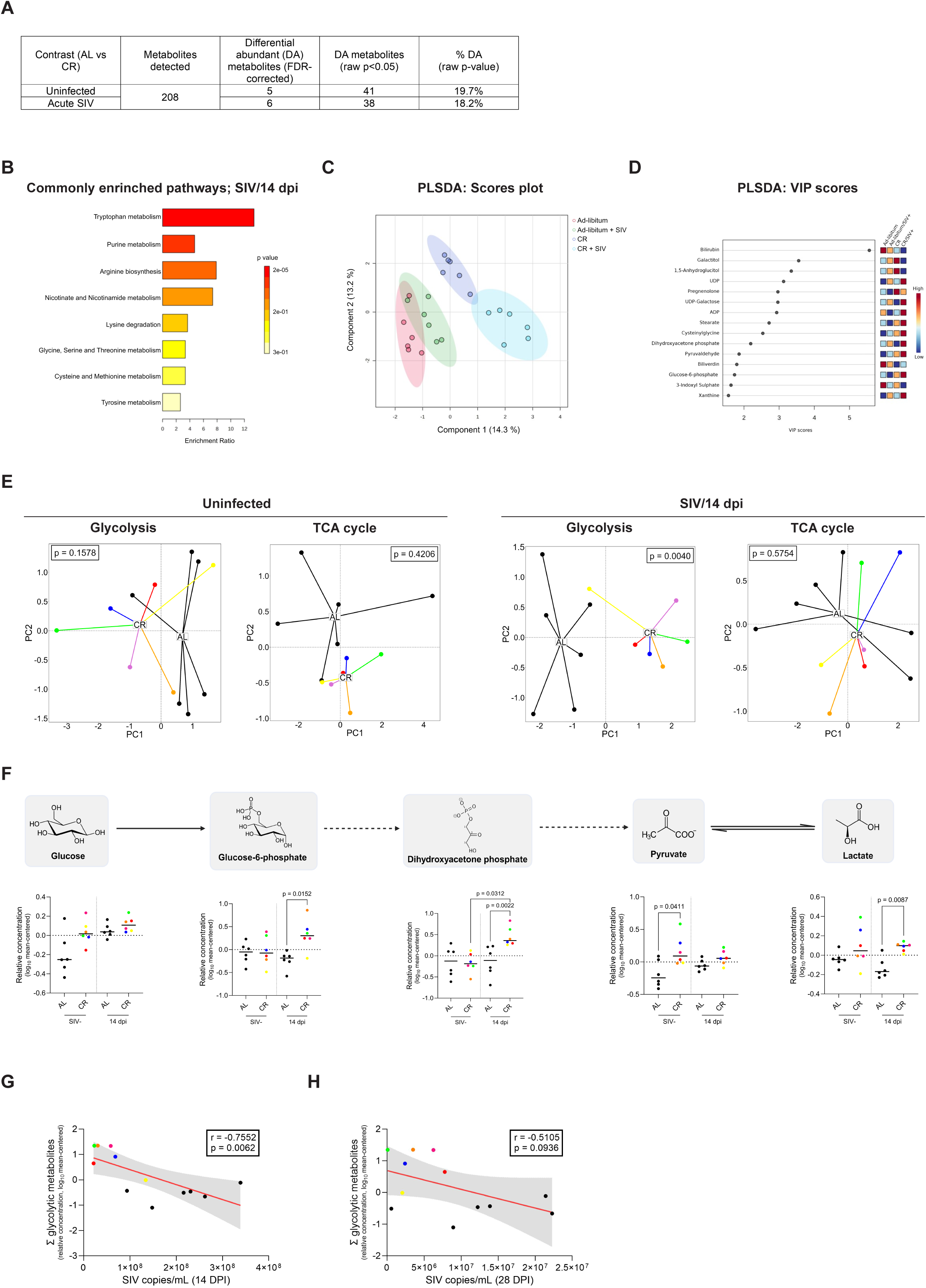
CR results in extensive alteration to metabolites in plasma before and after SIV. **(A)** Table depicting the differential abundance of plasma metabolites in AL (n=6) and CR (n=6) animals quantified through LC-MS at 14 dpi, showing numbers of differentially abundant (DA) metabolites that were significant by raw p and after correction for false discovery rate (Bonferroni post hoc test). **(B)** Bar chart identifying enriched metabolic pathways that are commonly differentially regulated at uninfected versus 14 dpi, generated via MetaboAnalyst 6.0. The enrichment ratio is calculated as the ratio of the observed number of hits within a specific metabolic pathway to the anticipated number of hits in that pathway. This analysis used KEGG as the reference database. The plasma metabolomic dataset was mean-centered and log10 normalized before any analysis. **(C)** Partial least square determinant analysis (PLS-DA) of all detected plasma metabolites in AL and CR animals before and after SIV infection, represented as a bi-plot of the top contributing components. The ellipses correspond to 95% confidence intervals of a normal distribution. **(D)** Variable importance in projection (VIP) scores of the top 15 plasma metabolites that significantly contribute to the observed variation. The colored boxes on the right represent the row-scaled abundance of the corresponding metabolite. **(E)** Principal component analysis of the plasma levels of key glycolytic and TCA cycle metabolites in AL and CR animals at baseline and 14 dpi. The metabolites used for analysis are as follows: glycolysis-glucose, G6P, DHAP, pyruvate, lactate & TCA cycle-citrate, *cis*-aconitate, α-KG, succinate, fumarate, and malate. **(F)** Relative concentrations (mean-centered log10 normalized) of plasma glucose, G6P, DHAP, pyruvate, and lactate in AL and CR animals at baseline and 14 dpi. **(G & H)** Two-sided spearman correlation analyses between the sum of relative concentrations of all glycolytic metabolites and plasma viral load at 14 dpi **(G)**, and 28 dpi **(H)**. **Statistical analyses: (E)** PERMANOVA (Permutational multivariate analysis of variance), **(F)** Mann-Whitney U test and Wilcoxon signed-rank test for inter- and intra-cohort comparisons, respectively, and **(G & H)** two-sided spearman correlations with confidence bands generated through linear regression. α=0.05.

We also employed supervised clustering of plasma metabolites by Partial Least Squares Discriminant Analysis (PLS-DA) before and 14 days after SIV, which allowed us to segregate metabolic profiles by both diet and SIV serostatus (**Fig. 3C**). Strikingly, the effect size of CR (red vs dark blue ellipse) on the plasma metabolome was significantly greater than the effect size of acute SIV (red vs green ellipse) (**Fig. 3C**), indicative of broad CR-dependent metabolic reprogramming. We then extracted the Variable Importance of Projection (VIP) scores to identify the specific metabolites that distinguished AL and CR feeding regimens (**Fig. 3D**). Notable among these was adenosine diphosphate (ADP), a key molecule involved in cellular energy transfer. The concentrations of ADP in plasma were significantly higher in CR animals at 14 dpi (**Fig. S6A**). suggesting that CR either increased the precursor pool of ATP or enhanced the activity of ATP-coupling metabolic pathways. Plasma concentrations of cyclic adenosine monophosphate (cAMP), an pleiotropic ATP metabolite that triggers the protein Kinase A-dependent breakdown of hepatic glycogen stores^57^, significantly declined with acute SIV in AL-fed animals but remained stable with CR (**Fig. S6A**). Other metabolites increased by CR included several associated with the processing of β-galactose into glucose-1-phosphate (**Fig. S6B)** ^58^, together suggesting that CR increased metabolic reactions that generated glycolytic substrates endogenously.

We next determined the impact of CR on metabolites involved in glycolysis and the TCA cycle, which support key antiviral immune processes including the clonal expansion and effector function of immune cells. Principal component analysis revealed that plasma concentrations of both glycolytic and TCA cycle metabolites were similar between dietary groups prior to SIV infection (**Fig. 3E**). During acute SIV however, glycolytic but not TCA-cycle metabolites markedly segregated CR from AL animals (**Fig. 3E, S7A**). Indeed, several glycolytic intermediates were highly elevated in the plasma of CR animals during acute SIV, including G6P, dihydroxyacetone phosphate (DHAP), and lactate (**Fig. 3H**). Importantly, the lactate/pyruvate ratio remained unchanged (**Fig. S7B**), indicating that pyruvate increased proportionally with lactate production and that overall glycolytic flux was enhanced by CR. When summing the relative abundances of all glycolytic metabolites in plasma at 14 dpi, we found that these levels were inversely correlated to plasma viremia at 14 dpi. (**Fig. 3G**). The levels of glycolytic metabolites at 14 dpi were not predictive of VL at 28 dpi however (**Fig. 3H**), indicating a robust but temporally limited association of glycolysis with SIV protection.

### CR enhances cellular glycolytic flux and promotes mTOR activation during acute SIV infection

To determine whether glycolytically skewed profiles in plasma of CR animals were reflective of cellular metabolic states, we compared metabolic phenotypes in PBMCs from each dietary regimen by Seahorse flux analysis before and 14 days after SIV infection. Overall, extracellular acidification (ECAR), a proxy for glycolysis, was elevated in the PBMCs of CR animals (**Fig. 4A**). Measures of basal glycolytic rate, glycolytic capacity, and glycolytic reserve, suggested to represent the amount of reserved energy available in a cell upon responding to stress^59^, were elevated in PBMCs of CR animals prior to SIV infection and tended to remain elevated during acute SIV infection (**Fig. 4B**).

**Figure 4.**
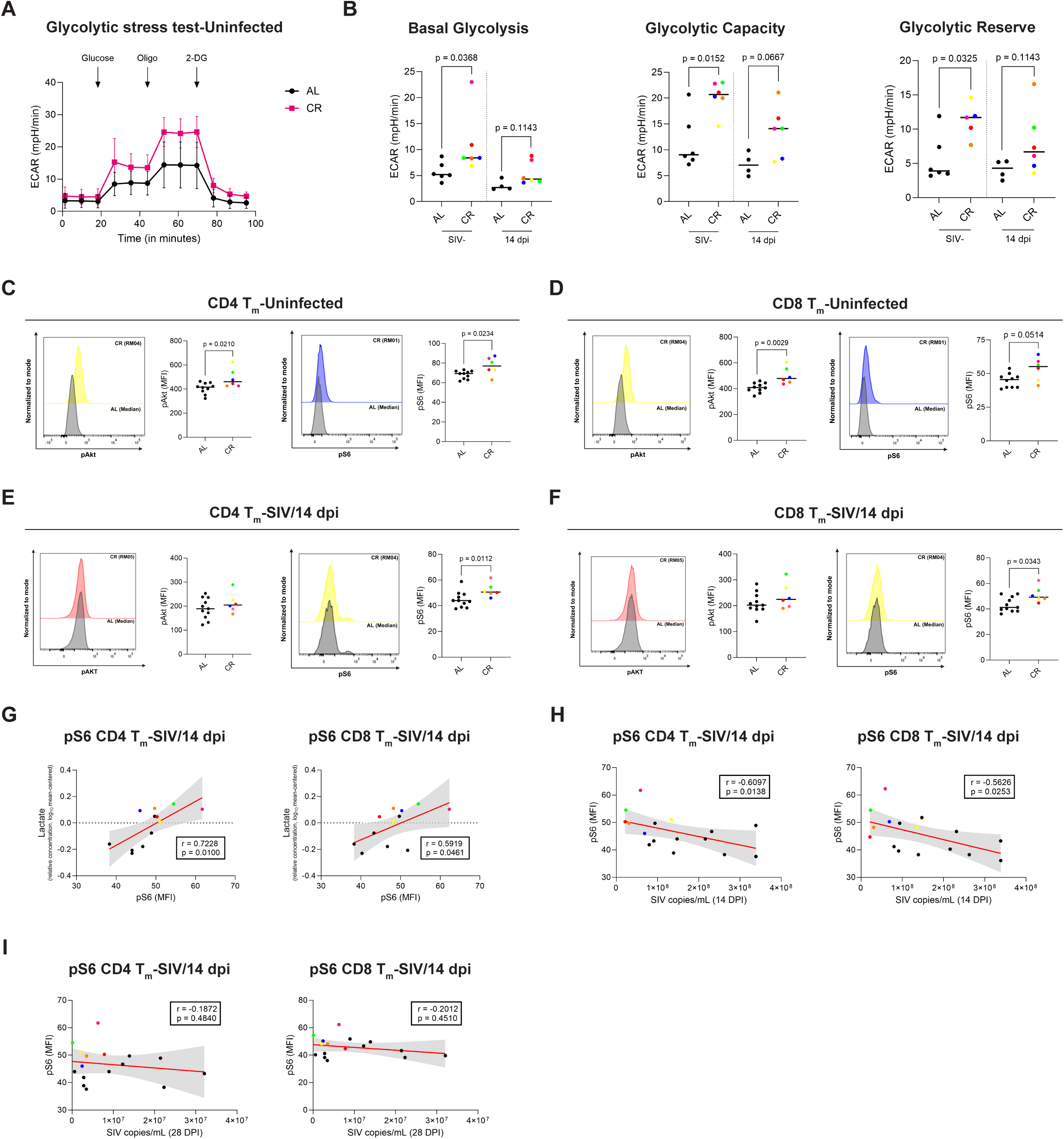
CR enhances cellular glycolytic flux and targets of mTORC activation in uninfected and acute SIV infected animals. **(A)** Metabolic profiling of PBMCs isolated from AL (AL, n=7; CR, n=6) and CR (AL, n=4, CR, n=6) animals at baseline using a Seahorse assay. The glycolytic pathway is assessed through real-time measurements of extracellular acidification rate (ECAR; mpH/min). **(B)** Indices of cellular glycolytic activity measured using Seahorse assay (basal glycolysis, glycolytic capacity & glycolytic reserve) in PBMCs isolated from AL and CR animals at baseline (AL, n=6; CR, n=6) and 14 dpi (AL, n=4, CR, n=6). Reduced data points in the AL group is due to lack of sufficient samples. **(C-F)** MFI levels of phospho-Akt (left) and pS6 staining (right) in circulating CD4^+^ total memory T cells of uninfected AL and CR animals **(C)** and 14 dpi **(E)** acquired via flow cytometry. MFI of pAkt (left) and pS6 staining (right) in circulating CD8^+^ total memory T cells of AL and CR animals at baseline **(D)** and 14 dpi **(F)**. Representative histograms of MFIs for each protein are shown next to their corresponding dot plots. **(G)** Two-sided spearman correlations of pS6 MFI in circulating CD4^+^ Tm and CD8^+^ Tm cells against relative concentrations of plasma lactate in AL and CR animals at 14 dpi. **(H)** Two-sided spearman correlations of pS6 MFI in circulating CD4^+^ Tm and CD8^+^ Tm cells in AL and CR animals at 14 dpi against their respective plasma viral loads (SIV copies/mL) at 14 dpi. **(I)** Two-sided spearman correlations of pS6 MFI in circulating CD4^+^ Tm and CD8^+^ Tm cells in AL and CR animals at 14 dpi against their respective plasma viral loads (SIV copies/mL) at 28 dpi. **Statistical analysis: (B-F)** Mann-Whitney U test and **(G-I)** Spearman non-parametric correlation with confidence bands generated through linear regression. α=0.05.

To further investigate cellular metabolic phenotypes associated with CR-mediated viremic protection at 14 dpi, we assessed the activation levels of Akt (pS473) and ribosomal protein S6 (pS235), both of which are phosphorylated downstream of the mammalian target of rapamycin (mTOR), a serine/threonine kinase that links environmental cues to glycolytic programs and the differentiation of effector T cells^60,61^. Prior to SIV, CR enhanced the phosphorylation levels of both Akt and S6 in circulating memory CD4⁺ and CD8⁺ T cells (**Fig. 4C, D**). Although we did not observe Akt phosphorylation to differ between groups during acute SIV (**Fig. 4E, F**), S6 phosphorylation remained elevated at 14 dpi in both CD4⁺ and CD8⁺ memory T cells and directly correlated with anaerobic production of lactate in plasma (**Fig. 4F, G**). Despite CR-mediated increases in pS6 activity, GLUT1, a facilitative glucose transporter in T cells typically linked to mTORC1 signaling^62^, was expressed at similar levels on CD4⁺ and CD8⁺ memory T cells in both AL and CR animals (**Fig. S7C**), indicating that activation of mTORC targets were independent of glucose uptake capabilities in circulating T cells. Consistent with trends observed for plasma glycolytic metabolite concentrations, pS6 activity in circulating CD4⁺ and CD8⁺ memory T cells were strongly associated with reduced viral load at 14 dpi but were not predictive of VL at 28 dpi (**Fig. 4H, I)**. Together, these findings indicate that glycolytically skewed plasma and cellular metabolic states are linked to early virologic protection, but this association diminishes as infection progresses.

### CR reduces immune activation in the GI tract and lowers systemic inflammation during ART

To understand the impact of CR on residual immune pathologies during ART and persistence of the SIV reservoir, beginning 28 dpi we administered a daily ART regimen of tenofovir (TDF), emtricitabine (FTC), and dolutegravir (DTG) by subcutaneous injection (**Fig. 2A**), which was sufficient to suppress viremia below detection limits in all animals by 11 months of continuous therapy (**Fig. S8A**). Initiation of ART under caloric- and time-restricted feeding was safe and well tolerated, and all animals successfully adhered to the CR diet up until the 11-month ART-suppressed study endpoint. The kinetics of viral decay following ART initiation did not differ significantly between the two cohorts (**Fig. S8B)**. Over the course of ART-treated SIV infection, both weights and body condition scores of CR animals remained lower than those of the AL-fed group (**Fig. 5A, B)**. Fasting blood glucose levels after 11 months of ART were comparable between dietary regimens (**Fig. 5C**). However, blood triglycerides were starkly reduced with CR across treated SIV (**Fig. 5D**), suggesting that CR improved the processing of lipids during ART.

**Figure 5.**
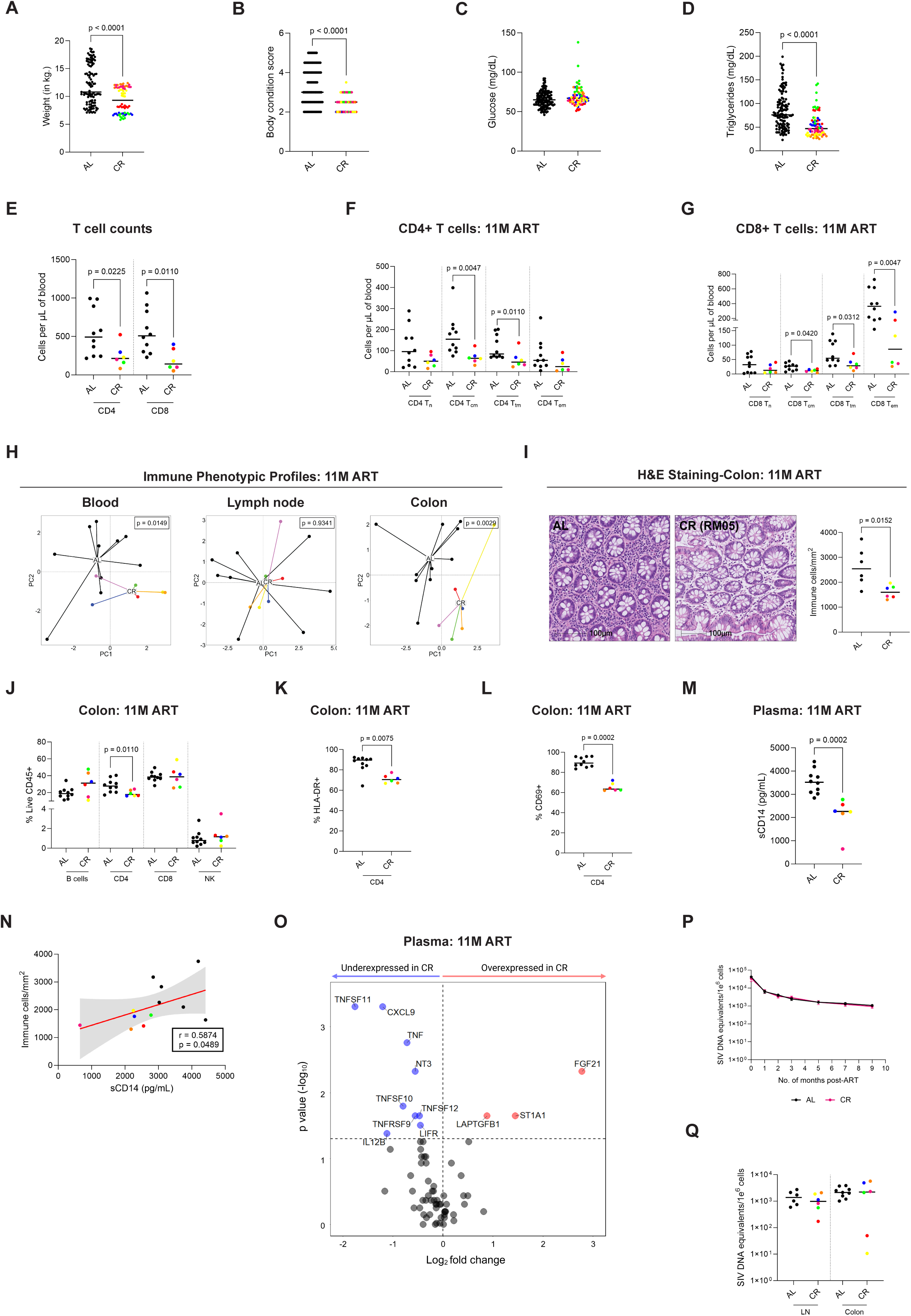
CR mitigates residual gastrointestinal dysfunction and inflammation during ART without impacting size of the SIV reservoir in blood and tissues. **(A & B)** Combined weight (in kg.) **(A)** and body condition scores **(B)** of AL and CR animals comprising all measurements throughout the entire duration of ART. **(C & D)** Combined plasma glucose **(C)** and triglyceride **(D)** (mg/dL) of AL and CR animals comprising all measurements throughout the entire duration of ART. **(E)** Absolute counts (cells/µL) of total CD4^+^ and CD8^+^ T cells in blood from AL and CR animals after 11 months of ART. Counts were calculated using total lymphocyte counts from complete blood chemistries and multiplied by frequencies of viable CD3^+^ CD4^+^ or CD8^+^ lymphocytes assessed via flow cytometry **(F & G)** Absolute counts (cells/µL) of CD4^+^ **(F)** and CD8^+^ T cell **(G)** maturation subsets in the blood of AL and CR animals at 11M ART. **(H)** Principal component analysis (PCA) of 15 immune phenotypic markers in distinct immune cell lineages (CD4^+^/CD8^+^ Tn, Tcm, Ttm, Tem, Treg (only CD4^+^), Tfh (only CD4^+^), NK cells (CD16^+^ & CD16^-^), and monocytes (classical, intermediate, & non-classical) in blood, lymph nodes, and colon at 11M ART, obtained via flow cytometry. Gating definitions were performed in accordance with Figure S2. **(I)** Representative H&E staining image of colon sections from one AL and CR animal (left). The graph on the right represents the quantification of immune cell infiltrates (immune cells/mm^2^) (right) at 11M ART. **(J)** Flow cytometric proportions of major immune cell subsets in the colon at 11M ART between AL and CR animals. **(K & L)** Percentage of total CD4^+^ T cells expressing HLA-DR **(K)** and CD69 **(L)** in the colon at 11M ART obtained via flow cytometry. **(M)** Quantification of soluble CD14 (sCD14) in the plasma at 11M ART through ELISA. Values were interpolated using a standard curve and expressed in pg/mL. **(N)** Correlation between plasma sCD14 (ELISA) and colon immune cells/mm^2^ (H&E) at 11M ART. **(O)** Volcano plot representing the mean log2 fold change of plasma cytokine and chemokine concentrations at 11M ART. red = over-expressed in CR, blue = under-expressed in CR, gray = nonsignificant. **(P)** Quantification of total SIV DNA in purified blood CD4^+^ T cells at ART initiation through 9 months of continuous ART. Data is represented as SIV DNA cell equivalents per million cells. **(Q)** Quantification of total SIV DNA in the LNs and colon at 11M ART. Data is represented as cell equivalents per million cells. **Statistical Analyses: (A-G, I-M, P-Q)** Mann-Whitney U-test, **(H)** PERMANOVA and **(N)** Spearman non-parametric correlation with confidence bands generated through linear regression. α=0.05.

We next determined the impact of dietary regulation on immune reconstitution by examining numbers of CD4⁺ and CD8⁺ T cells in blood, the former of which typically rebound with ART initiation. Numbers of both circulating CD4⁺ and CD8⁺ T cells were unexpectedly lower in CR animals after 11 months of ART (**Fig. 5E**), which were driven specifically by the reduction of central and transitional memory T cells in the CD4⁺ compartment (**Fig. 5F**), and reductions of all memory subtypes (central, transitional, effector) in the CD8⁺ T cell compartment, with effector representing the starkest reduction (**Fig. 5G**). Lymphopenia was unlikely to be driven by increased rates of cell death under CR, as intracellular expression of the pro-survival molecule Bcl-2 was comparable between dietary regimens (**Fig. S8C, D**). Moreover, proportions of active caspase-3-expressing T cells were elevated in CD8⁺ T cells of ART-suppressed AL-fed animals despite their higher numbers in circulation (**Fig. S8F**). CR is well known across species to promote the retention of blood monocytes and T cells within the bone marrow (BM)^35,63^. Although memory CD8⁺ T cells were reduced in blood, we observed their frequencies were increased in BM during ART (**Fig. S8G)**. We also observed that frequencies of intermediate monocytes in CR animals were likewise increased at this tissue site during ART (**Fig. S8H**), suggesting retention of T cells and monocytes in the bone marrow that is a known ‘safe haven’ for memory T cell homeostasis^35,63^. Principal component analysis of 15 immune phenotypic markers on CD4^+^ and CD8^+^ T cells, NKG2A^+^ NK cells, and myeloid cells across tissues revealed that the anatomic site exhibiting the broadest immunologic response to CR was the colon (**Fig. 5H**). To examine this further, we performed Hematoxylin & Eosin (H&E) and Myeloperoxidase-1 (MPO-1) staining in the gut to quantify lymphocytes and granulocytes, respectively. H&E staining revealed ongoing diffuse lymphocytic infiltration throughout the lamina propria in ART-suppressed AL-fed animals, which was markedly reduced by CR (**Fig. 5H**). Myeloid cell numbers in the colon remained comparable between dietary regimens in ART-suppressed animals (**Fig. S8I**). In assessing the potential lineage of infiltrated/expanded leukocytes in the colon by flow cytometric analysis, only CD4⁺ T cell percentages were distinctly elevated in the AL-fed colon during ART (**Fig. 5J**). Colonic CD4⁺ T cells were significantly more activated, as highlighted by their surface expression of both HLA-DR and CD69 (**Fig. 5K, L**). This trait was unique to the colon in AL-fed animals and not observed in the blood or lymphoid tissues (**Fig. S8J, K**). Phenotypically, the infiltrating CD4⁺ T cells in the AL colon were significantly more Th1-like by exhibiting higher CXCR3 expression (**Fig. S8L**). In contrast, CD4⁺ T cells in the CR-colon were proportionally more skewed towards CXCR5^+^ PD-1^+^ T_fh_ cells and tended to be enriched for a CD25^+^ Foxp3^+^ T regulatory cell phenotype (**Fig. S8M, N)**, suggesting a more tolerogenic state.

Given the known link between GI dysfunction and residual inflammation during ART, we assessed the concentrations of plasma soluble CD14 (sCD14), a marker of monocyte exposure to Lipopolysaccharide (LPS) that is highly predictive of earlier mortality in ART-suppressed PWH^64^. CR markedly reduced plasma sCD14 levels after 11 months of ART (**Fig. 5M**). We observed a direct relationship between sCD14 levels in plasma and densities of activated leukocytes in the colon (**Fig. 5N**). Because a damaged gut in HIV-1/SIV can sustain systemic inflammation, we next determined whether CR limited residual inflammation on ART. A panel containing more than 60 soluble inflammatory markers revealed a significant downregulation of the plasma inflammatory profile with CR, including a reduction of the Type-II Interferon-induced protein CXCL9, and several NF-κB responsive cytokines of the Tumor Necrosis Factor (TNF) family (TNF, TNFSF9-12) (**Fig. 5O**). Plasma concentrations of the TGFβ precursor protein LAPTGFB1 and fibro-blast growth factor 21 (FGF21) were soluble factors that were upregulated with CR during ART suppression (**Fig. 5O**).

Residual inflammation during ART exists concurrently with the persistence of viral reservoirs, and some groups have noted directed associations with certain inflammatory markers and levels of HIV-1 DNA in blood^65,66^. We thus measured ART-mediated decay dynamics of the SIV reservoir between AL and CR groups that exhibited diverging levels of inflammation. Despite a dampened systemic inflammatory profile under CR, the decay dynamics of total SIV DNA in blood CD4⁺ T cells following ART initiation were highly similar to those of AL-fed animals (**Fig. 5P**). We also did not observe differences in total SIV DNA within pLN CD4⁺ T cells or bulk mononuclear cells in the colon between dietary regimens after 11 months of ART (**Fig. 5Q**). The fact that many systemic inflammatory mediators were mitigated by CR, but SIV DNA levels were unchanged suggest that in this particular setting, dynamics of the reservoir during ART were not influenced by inflammation or gut mucosal immune dysfunction.

### CR increases relative concentrations of bile acids and TCA-cycle metabolites in plasma during ART suppression

Examining plasma metabolomes at 10 months of ART revealed complete segregation by dietary regimen (**Fig. 6A**). The unique metabolic profile in plasma of ART-suppressed CR animals was over-represented by several bile acids (**Fig. 6B**). Specifically, concentrations of both primary (cholate) and secondary bile acids (tauromuricholate, taurodeoxycholate, glycochenodeoxycholate) were elevated in plasmas of CR animals during ART-suppression (**Fig. 6C**). Elevated bile acid concentrations in plasma can be an indicator of liver damage (refs). The CR-dependent increases in plasma bile acids however were independent of any difference to liver enzymes in the serum chemistry panel (ALT/AST/bilirubin) (**Fig. S9A-C)**, suggesting rather an overall improvement of bile acid homeostasis with CR.

**Figure 6.**
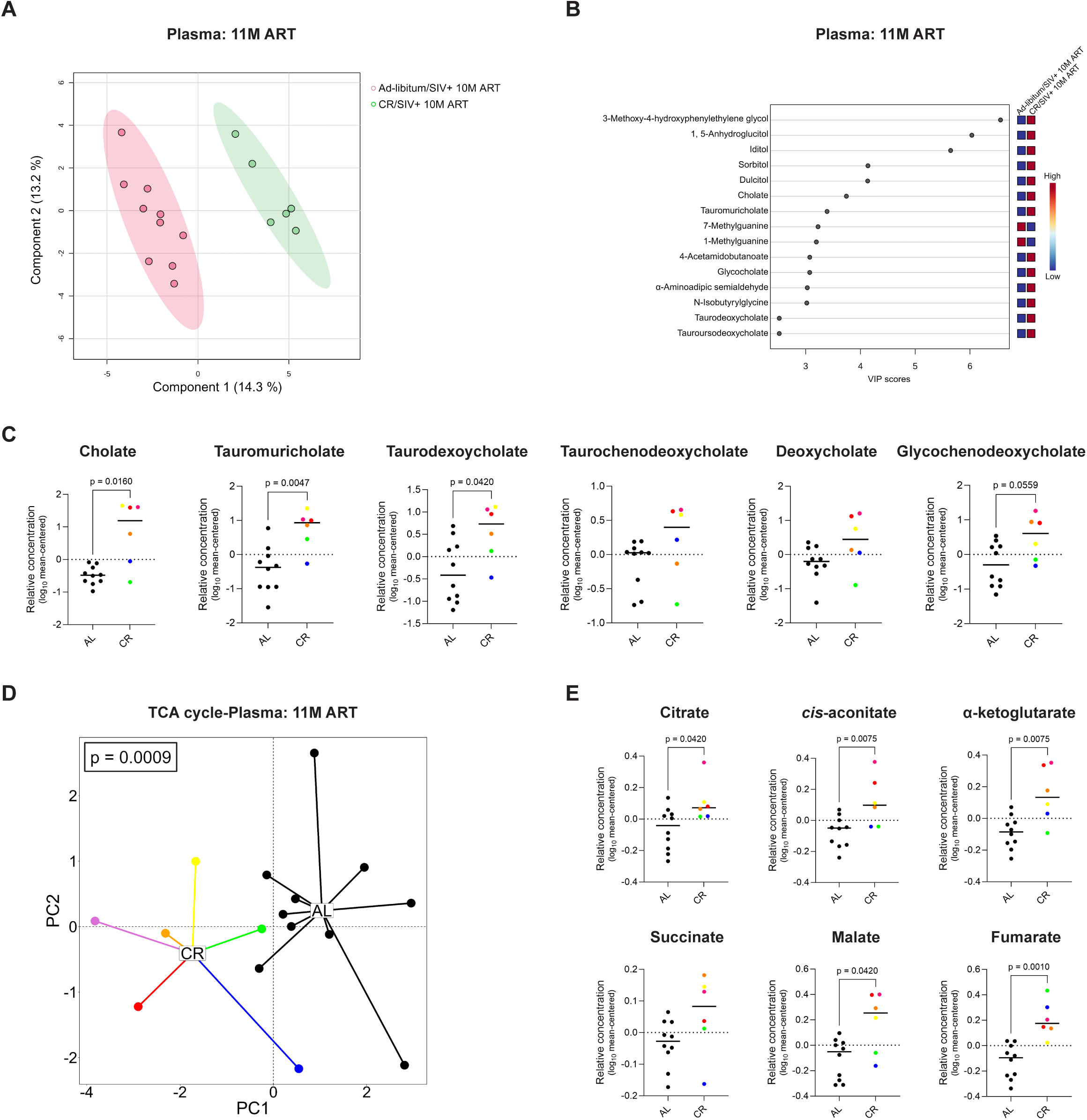
Plasma concentrations of bile acids and TCA-cycle intermediates are increased with CR during ART. **(A)** PLS-DA analysis of all detected plasma metabolites after 11 months of ART in AL and CR animals, represented as a bi-plot of the top contributing components. The ellipses correspond to 95% confidence intervals of a normal distribution. **(B)** Variable importance in projection (VIP) scores of the top 15 plasma metabolites that significantly contribute to the observed variation. The colored boxes on the right represent the row-scaled abundance of the corresponding metabolite. **(C)** Relative concentrations (mean-centered log10 normalized) of plasma bile salts in AL and CR animals at 11M ART. **(D & E)** Principal component analysis (PCA) of key TCA cycle intermediates in the plasma **(D)** and their relative concentrations **(E)** in AL and CR animals at 11M ART. **Statistical analyses: (C & E):** Mann-Whitney U test and **(D):** PERMANOVA. α=0.05.

Given the prominent plasma glycolytic profile induced by CR during acute phase viremia (**Fig. 3E, F**), we assessed whether this phenotype persisted after 11 months of ART. When all detected glycolytic metabolites were analyzed as a principal component, CR animals remained distinct from their AL counterparts, though less significantly so than during acute-phase untreated infection (**Fig. S10A**). Moreover, the metabolites driving segregation during ART suppression differed from those in untreated SIV, with CR-dependent increases in glucose and pyruvate being the primary contributors (**Fig. S10B**). Unlike during untreated SIV infection, concentrations of TCA-cycle metabolites were highly distinct between dietary regimens after 11 months of ART (**Fig. 6D**). All TCA cycle metabolites, except succinate, which was also notably increased, were significantly elevated in the plasma of ART-suppressed CR animals (**Fig. 6E**). Taken together, these data suggest that CR induced a phase-dependent metabolic programming during SIV infection, characterized by glycolytic dominance during acute infection, while increasing TCA cycle flux during ART-suppressed chronic infection. Both of these metabolic profiles occurred in the context of a more limited inflammatory state during untreated and treated SIV.

## DISCUSSION

Immunometabolic pathways are co-opted by HIV-1 to sustain each step of the viral replication lifecycle^15,17^. They also underpin effective antiviral immunity^67^, and their dysfunction in HIV-1 contributes to hallmark pathologies including immune exhaustion and chronic inflammation^22^. Despite the broad involvement of immunometabolism in HIV-1 infection, it is not clear how diet, a master regulator of metabolic behavior, can influence the course of infectious diseases in higher-order mammals. Moreover, distinct metabolic phenotypes that are associated with favorable outcomes in the setting of HIV-1 are not well described. In this study, we safely regulated dietary intake with the goal of manipulating metabolism to improve virologic and immunologic outcomes in SIV infection. We found that 4 months of CR conditioning reduced viral loads and evoked a more constrained type-I Interferon response when compared to that of AL-fed animals. During ART, CR lowered the densities of activated leukocytes in the colon and mitigated several soluble innate immune mediators whose levels in plasma predict mortality in PWH^4^. Overall, the data highlights that safely manipulating the timing and quantity of caloric intake can improve several clinically meaningful metrics of a primate lentiviral infection.

Arguably the most striking observation of this study was that CR reduced acute phase SIV viral loads. Lower viremia was better predictive of a more limited pool of target cells in the CR colon prior to SIV, a major early site of viral replication^2,45,46^. Concurrently, lower viral loads in CR animals were met with a more restrained induction of type-I interferon in plasma. These levels correlated inversely with the cycling of memory CD8^+^ T cells in lymphoid tissues, suggesting the absence of a strong anti-proliferative effect that is a well-known feature of high exposure to type-I interferon^55,56^. Indeed, while we were unable to directly block inflammatory molecules by mechanistic studies, our data suggest that the enhanced cycling of CD8^+^ T cells in CR animals was a consequence rather than a cause of lower plasma SIV, as neither the viral growth nor decay rate of plasma virus differed between dietary regimens. Moreover, barcode sequencing of individual replicating plasma SIV strains revealed comparable community structure between groups, arguing against a selective pruning of distinct viral lineages by the immune system.

An additional notable finding of this study was that during acute SIV, CR induced a robust up-regulation of glycolytic metabolites whose concentrations in plasma were inversely correlated to peak VL at 14 dpi. Our data supports the notion that CR acutely magnifies the ‘Warburg effect,’ characterized by the preferential utilization of glycolysis despite oxygen-replete conditions. These findings are remarkably concordant with protective metabolic signatures in *M. tuberculosis*-infected mice on a CR diet^33^ and indicate that glycolytic skewing of immunity upon infectious challenge is a feature of CR that transcends both species and pathogen. While we were not able to interrogate this signature in mechanistic detail, glycolytic pathways could have supported the early clonal expansion, differentiation, and effector functions of antiviral immune cells, consistent with extensive evidence from murine models^9,14,68,69^.

To our knowledge, these data are the first to associate a protective role to glycolysis in a primate lentiviral infection. Critically, it is important to point out however that CR-dependent viral load reductions were transient and were not sustained out to 28 dpi. This may allude to the relative limits of a glycolytically dominated environment in viral infections. Indeed, glycolytic pathways can support the HIV-1 replication lifecycle *in vitro* (ref).

There is also a large body of evidence that in chronic HIV-1/SIV infection settings, sustained and monotypic glycolytic signatures are linked to chronic inflammation, immune activation, and immune exhaustion in both untreated and ART-suppressed PWH^22,70,71^ ^67^. The dichotomous nature of glycolysis is highlighted in our own studies. For example, we found that plasma levels of the glycolytic end product lactate were highly elevated in CR animals and were linked to lower viral loads in early acute SIV. Chronic lactate exposure however is detrimental to a variety of immunological processes and can potentiate inflammation, and buildup of lactate may have contributed to the ultimate waning of virologic control in our studies^72–76^. These data highlight that the immunologic benefits of a glycolytically dominated response to viral infections can be robust but ultimately short-lived.

In line with this contextual framework, we show here that the metabolic phenotype-In the plasma of CR animals transitioned to one enriched for TCA-cycle metabolites during fully suppressive ART. Importantly, high concentrations of these intermediates were concurrent with reductions in particular soluble mediators of innate immune activation that predict mortality in ART-suppressed PWH^4^. While the nature of a TCA-cycle dominated phenotype to anti-inflammatory states in HIV-1/SIV is not clear, one potential link could be the ability of this metabolic pathway to sustain homeostasis of the GI tract. Gut colonocytes, which comprise 80-90% of the colonic epithelium^77^, are highly oxidative^78^, and glutamine and its conversion to α-KG is a dominant energy pathway that sustains enterocyte proliferation and integrity of the gut epithelial barrier. The capacity of the GI tract to maintain metabolic flux through the TCA cycle may thus be critical to fully support repair of a damaged gut upon ART initiation.

An observation extending beyond immunometabolism with relevance to HIV-1 cure studies was that relative to AL-fed controls, the amount of total SIV DNA in blood and tissues of CR animals was not different despite a significantly reduced inflammatory state. The forces driving both viral persistence and residual inflammation exist concurrently during long-term ART, and at least two studies have observed particular inflammatory mediators in plasma to correlate directly with intact HIV-1 DNA^65,66^. It is important to note, however, that correlation does not imply causality. In our study, the particular inflammatory profile mitigated under a CR diet consisted mainly of TNF superfamily molecules and soluble interferon-stimulated proteins (CXCL9). While we cannot rule out that an inflammatory milieu unique to this signature could indeed actively replenish the reservoir, our data suggests that this inflammatory profile does not drive the persistence of reservoir cells. On the contrary, there is more evidence to suggest that factors influencing the proliferation, survival, and cellular death rates of CD4⁺ T cells are more determinative in shaping overall reservoir dynamics^79–82^. Inflammatory mediators that work in tandem with these physiological processes could, in theory, influence reservoir dynamics. For example, TNF-enriched signatures in expanded CD4⁺ T cell clones from ART-suppressed PWH have been observed to correlate directly with the size of that clone under resting and antigen-stimulated conditions^83^. In mice, TNF can also enhance the *in vivo* clonal expansion of CD4⁺ T cells^84^. It is thus possible that these inflammatory cytokines could serve as co-stimulatory signals to augment the antigen-induced clonal expansion of reservoir cells. Nevertheless, our data suggests that, at least in the well-established macaque model of SIV infection, blockade of inflammatory mediators over 11 months of ART would not meaningfully reduce the size of the reservoir. In line with this finding, several small-molecules tested in pre-clinical and clinical settings that reverse HIV-1/SIV latency do so by inducing systemic NF-κB or Type-I Interferon responses, yet these are not met with reservoir expansion^85,86^, and in some cases slightly reduce levels of intact HIV-1 DNA^85^.

### Limitations of the study

First, we note in our study that CR was implemented within a time-restricted feeding window. While we observed that the 2-month TR conditioning phase prior to implementation of CR did not result in changes to blood glucose, lipid levels, or T cell activation state, we were not able to assess the variable of TR alone on metabolic, virologic, and immunologic dynamics during SIV. One animal in our cohort, RM03, underwent calorie but not time restricted feeding, and responses to SIV in this animal did not appear to dramatically diverge from those undergoing CR in a TR feeding window. While anecdotal, this observation is consistent with time restricted feeding regimens having a comparably more muted impact on host physiology than CR^37,87^. Second, an ART treatment interruption (ATI) could not be performed in this study, as control AL-fed animals were required to remain on ART beyond the 10-month CR endpoint due to participation in a concurrent ancillary study. We were thus not able to assess the impact of CR on post-ATI viral rebound. Last, we emphasize that CR in this study was established preventively, before SIV infection. It is currently unknown whether CR would be equally efficacious in a therapeutic setting under the context of pre-existing tissue damage during ART. Short-term CR, or reducing caloric intake pharmacologically by mimicking satiety with incretin mimetics^88^, may represent a more clinically relevant extension to this study.

## METHOD DETAILS

### Animals & Experimental Model

This study included 17 adult, non-obese, male and female Indian-origin rhesus macaques divided into two age-, sex-, and weight-matched cohorts. All macaques were bred and housed at Tulane National Primate Research Center (TNPRC) in compliance with the NRC guide for the Care and Use of Laboratory Animals and the Animal Welfare Act. The Tulane National Primate Research Center (TNPRC) is a fully accredited institution by AAALAC International, Animal Welfare Assurance No. A3180-01. All experimental procedures performed as part of the study were approved by the Institutional Animal Care and Use Committee (IACUC) of Tulane University.

Except for the initial *ad libitum* period, all animals were pair-housed indoors under well-ventilated and climate-controlled conditions. Throughout the study, all procedures were carried out under general anesthesia induced by intramuscular (i.m.) injection of Telazol^®^ (tiletamine hydrochloride and zolazepam hydrochloride; Zoetis, Parsippany-Troy Hills, NJ) at a dose of 3-8 mg/kg. Animals also received prophylactic and post-procedural analgesics to minimize the pain and distress experienced as a result of any clinical and/or surgical procedures. Following the end of the study, all animals were euthanized in accordance with IACUC and American Veterinary Medical Association (AVMA) guidelines.

### SIV infection and ART initiation

All animals were intravenously (i.v.) infected with 5000 IU/mL of genetically barcoded SIVmac239M virus ^89^. The SIVmac239M inoculum was a kind gift from Dr. Brandon Keele at Frederick National Laboratory for Cancer Research, Maryland, USA. At 28 dpi, all animals regardless of their experimental status were started on combined antiretroviral therapy (cART) composed of a regimen of three antiviral drugs: TDF (Tenofovir disoproxil fumarate; ∼5 mg/kg), FTC (Emtricitabine; ∼38 mg/kg), and DTG (Dolutegravir; ∼2.5 mg/kg) dissolved in distilled water containing 15% w/v KLEPTOSE^®^ HPB-LB Parenteral Grade carrier (Roquette Frères, Lestrem, France). This cART cocktail was administered subcutaneously every day at a dose of 1 mL/kg.

### Intravenous glucose tolerance test

The intravenous glucose tolerance test (IVGTT) was conducted to assess glucose metabolism and insulin sensitivity. A 50% (w/v) dextrose solution was infused intravenously over approximately 30 seconds. Capillary blood glucose levels were measured using a glucometer at baseline (prior to dextrose infusion) and at the following time points: 3-, 5-, 7-, 10-, 15-, 20-, and 30-minutes post-dextrose administration. The volume of the dextrose solution administered was calculated based on the most recent body weight of the animal, with the dosage being maintained at ∼0.33 mL/kg of body weight. Capillary blood glucose samples were collected via finger, ear, or tail stick at the specified time points, and by calculating the area under the curve (AUC) using the trapezoidal method to assess overall glucose tolerance^90^.

### Blood and tissue processing

All blood for the study was collected using EDTA-containing S-Monovette^®^ tubes (Sarstedt Inc., Newton, NC). Plasma isolation was carried out via centrifugation at 900 x g for 15 minutes, following which PBMCs were isolated via traditional density-gradient cell isolation using Ficoll-Paque^TM^ PLUS (Cytiva Life Sciences, Marlborough, MA) using the manufacturer’s instructions. Lymph nodes and colon pinches were homogenized through mechanical disruption using gentleMACS^TM^ Dissociator (Miltenyi Biotec, Gaithersburg, MD) and filtered through 100 μm cell strainers. All cells were cryopreserved in liquid nitrogen in a freezing medium consisting of fetal bovine serum (FBS) (Corning Inc., Corning, NY) with 10% v/v dimethyl sulfoxide (DMSO) (EMD Millipore, Burlington, MA).

### Complete blood counts and chemistries

Hematological analyses consisting of complete blood counts (CBC) and standard chem-20/SMAC-20 screenings were performed using ∼1 mL each of clotted and EDTA-treated blood, respectively, by our in-house clinical assay services core at TNPRC. This was done using the Sysmex NX-V-1000 Hematology Analyzer (Sysmex Corporation, Hyogo, Japan).

### SIV plasma viral load quantification

Plasma viral load quantification was carried out by the Pathogen Detection and Quantification Core (PDQC) of TNPRC as described by C J Monjure et al^91^. Briefly, SIV target cDNA and exogenous control cDNA were assayed in duplicate using the QuantStudio^TM^ 12K Flex system (Thermo Fisher Scientific, Waltham, MA). The cycling program consisted of 40 cycles at 95 °C for 15 seconds and 60 °C for 1 minute. The sequences of primers and probes used during PCR amplification are as follows: Forward 5′- AGGCTG-CAGATTGGGACTTG-3′, Reverse 5′-TGATCCTGACGGCTCCCTAA-3′, and Probe 5’- FAM-ACCCACAACCAGCTCCACAACAAGGAC-IABKFQ-3′. The viral decay rate was computed through non-linear regression and curve fitting to an exponential one-phase decay model. It is defined as the rate constant λ in the equation: Y = (Y_0_ - Plateau)*e^(-λ*𝜏𝜏)^ + Plateau, where 𝜏𝜏 denotes time and plateau denotes the LOD.

### Cell-associated (CA)-SIV DNA quantification

Approximately 1 – 1.5 x 10^6^ CD4⁺ T cells were immuno-magnetically separated from bulk mononuclear cell suspensions by positive selection using MACS^®^ MicroBeads conjugated to a monoclonal anti-human CD4 antibody as per the protocol (Miltenyi Biotec, Gaithersburg, MD). Separated CD4⁺ T cells or colonic cell suspensions were lysed with Buffer RLT Plus (Qiagen, Hilden, Germany) and 10% β-mercaptoethanol (Millipore Sigma, Burlington, MA), and DNA was subsequently purified using the AllPrep DNA spin column kit. The CA-SIV DNA quantification was performed employing a standard quantitative PCR protocol and TaqMan^TM^ Fast Advanced Master Mix (Applied Biosystems, Thermo Fisher Scientific, Waltham, MA). The PCR conditions included an initial step of 95 °C for 2 minutes for template denaturation (Thermo Fisher Scientific, Waltham, MA), followed by 40 cycles of 95 °C for 30 seconds and 60 °C for 60 seconds. The primers (and their respective reaction concentrations) used were as previously published and as follows^92^: *sGAGF* 5′- GTCTGCGTCATCTGGTGCATTC-3′ (600 nM), *sGAGR* 5′-CAC-TAGGTGTCTCTGCACTATCTGTTTTG-3′ (600 nM), *sGAGPr* 5′-FAM-CTTCCTCAG-TKTGTGTTTCACTTTCTCTTCTGCG-BHQ1-3′ (100 nM), *CCR5F* 5′-CCAGAA-GAGCTGCGACATCC-3′ (100 nM), *CCR5R* 5′-GTTAAGGCTTTTACTCATCTCAGAA-GCTAAC-3′ (100 nM) and *CCR5Pr* 5′-CalRed610-TTCCCCTACAA- GAAACTCTCCCCGGTAAGTA-BHQ2-3′ (100 nM). All reactions were quantified on a QuantStudio™ 6 Pro Real-Time PCR System and expressed as cell equivalents per million cells.

### SIVmac239M barcode sequencing and analysis

Viral RNA was isolated from plasma using the QIAamp Viral RNA Mini Kit (Qiagen, Hilden, Germany) according to the manufacturer’s instructions. Complementary DNA (cDNA) was synthesized using SuperScript IV reverse transcriptase (Invitrogen, Waltham, MA) with an SIV-specific reverse primer (Vpr.cDNA3; 5′-CAG GTT GGC CGA TTC TGG AGT GGA TGC-3′). cDNA was quantified by quantitative real-time PCR (qRT-PCR) using primers VpxF1 (5′-CTA GGG GAA GGA CAT GGG GCA GG-3′) and VprR1 (5′-CCA GAA CCT CCA CTA CCC ATT CAT C-3′) in combination with a labeled probe (5′-ACC TCC AGA AAA TGA AGG ACC ACA AAG GG-3′).

For next-generation sequencing, PCR amplification was performed using VpxF1 and VprR1 primers appended with Illumina P5 or P7 adaptor sequences and unique 8-nucleotide index barcodes to allow multiplexing. PCR amplification was carried out using Platinum High Fidelity Taq DNA Polymerase (Thermo Fisher Scientific, Waltham, MA). Indexed amplicons were pooled and sequenced on an Illumina MiSeq platform. Sequence processing and barcode analysis were performed as previously described^89,93^.

Barcode sequencing data were summarized using complementary measures of clonal diversity and inequality. The number of unique viral clones was used as a measure of barcode richness, reflecting the total number of distinct viral lineages detected in each sample. The Shannon diversity index was calculated to capture both barcode richness and the relative evenness of lineage abundances. The Gini diversity coefficient was used to quantify inequality in barcode distributions, with higher values indicating increased dominance by a limited number of viral clones. All diversity and clonal structure analyses were performed in RStudio using custom scripts.

### Immune phenotyping by flow cytometry

Cells were washed and counted using the Cellometer Auto 2000 (Nexcelcom Bioscience, Lawrence, MA). A minimum of 1.5 x 10^6^ cells were aliquoted and stained using pre-optimized concentrations of necessary antibodies (see **REAGENT RESOURCES TABLE**). Samples were stained at 37 °C for 15 minutes with the live/dead amine-reactive dye and then for another 20 minutes with surface antibodies. Permeabilization for intra-cellular staining was carried out using 500 µL of 1x eBioscience^TM^ Fixation/Permeabilization Concentrate diluted 1:4 using eBioscience^TM^ Fixation/Permeabilization Diluent (Invitrogen, Waltham, MA) by incubating surface-stained cells at 4 °C for 1 hour. This was followed by staining for intracellular markers at 4 °C for 30 minutes. All washes post-permeabilization were carried out using 1x eBioscience^TM^ Permeabilization buffer (diluted in sterile DPBS) at 900 x g for 5 mins. Following the final wash, cells were fixed with a 1% paraformaldehyde (PFA) solution before acquisition. The stained samples were acquired on a BD FACSymphony^TM^ A5 cytometer, and data analysis was performed using Flow-Jo^TM^ (v 10.10.0).

### Phospho-Akt and S6 flow cytometry

Phospho-staining for phosphorylated Akt and ribosomal protein S6 (or simple S6) was carried out using an optimized 7-color antibody panel. For this purpose, an appropriate no. of cells were aliquoted into FACS tubes, stained first with 2 µL of the live/dead stain and appropriate surface antibodies at 37 °C for 20 minutes. Cells were then washed at 900 x g for 5 minutes using sterile DPBS and fixed with 500 μL of BD Cytofix/Cytoperm Fixation/Permeabilization kit (BD Biosciences, Franklin Lakes, NJ) at 37 °C for 15 minutes. After this, cells were washed and resuspended in 200 µL of BD Phosflow^TM^ Perm Buffer III (BD Biosciences, Franklin Lakes, NJ), followed by incubation at 4 °C for 30 minutes. Cells were then washed and stained with intracellular antibodies (anti-Akt and anti-S6) and placed at 4 °C for 20 minutes. This was followed by a repeat wash and then fixation in 300 µL of 1% PFA for at least 2 hours. The fixed samples were acquired on an LSRFortessa™. Akt and S6 staining intensities were analyzed using FlowJo and reported as median fluorescence intensity (MFI).

### Live PBMC Seahorse Assay

Metabolic stress responses in live PBMCs were captured through the Agilent Seahorse Glycolytic Stress test (GST) using the Agilent XFe24/96 Seahorse Analyzer (Agilent Technologies, Santa Clara, CA). Before beginning the assay, sensor cartridges supplied with the experimental kit were hydrated using Seahorse XF Calibrant Solution (Agilent Technologies, Santa Clara, CA) overnight. The following day, 5x10^5^ freshly isolated PBMCs resuspended in assay medium are seeded into each well of the 24-well plate, with one well per row containing only plain assay medium for background correction. The assay medium for GST is Seahorse XF RPMI medium supplemented with 2 mM glutamine. The cells were rested for a minimum of 30 minutes (not to exceed 1 hour) in a CO2-free incubator at 37 °C. During GST, extracellular acidification rate (ECAR; mpH/min) and oxygen consumption rate (OCR; pmol/min) were measured in real time by sequentially exposing the cells to 100 mM glucose, 10 µM oligomycin, and 500 mM 2-deoxyglucose (2-DG). The cellular glycolytic activity was estimated from the ECAR profile as follows: basal glycolysis represents the ECAR difference from baseline and the addition of glucose, glycolytic capacity is the maximum ECAR measurement captured following oligomycin, and glycolytic reserve is defined as the difference in oligomycin ECAR and 2-DG ECAR. All samples were run in 5 replicates, and data points are expressed as median ± SEM.

### Olink^®^ cytokine and chemokine quantification

Proteins were measured using the Olink^®^ Target 96 panel (Olink Proteomics AB, Uppsala, Sweden) by our in-house High Containment Research Performance Core (HCRPC), following the manufacturer’s instructions. The Proximity Extension Assay (PEA) technology used for the Olink protocol has been well described by Assarsson et al. and enables 92 analytes to be analyzed simultaneously using 1 µL of each sample ^94^. In brief, pairs of oligonucleotide-labeled antibody probes bind to their targeted protein, and if the two probes are brought in close proximity, the oligonucleotides hybridize in a pair-wise manner. The addition of a DNA polymerase leads to a proximity-dependent DNA polymerization event, generating a unique PCR target sequence that is subsequently detected and quantified using the Fluidigm^®^ BioMark HD microfluidic real-time PCR instrument. The resulting Ct-data underwent comprehensive quality control and normalization using a set of internal and external controls. Four internal controls monitored different steps of the PEA process: two non-human proteins with matching antibody-probes functioned as incubation/immune-controls, an IgG antibody with two matching probes attached served as an extension control, and a complete double-stranded amplicon functioned as a detection control. External controls included a triplicate of negative controls used to calculate the limit of detection (LOD) and a triplicate of interplate controls (IPCs) used for normalization. The final assay read-out is presented in Normalized Protein eXpression (NPX) values, which is an arbitrary unit on a log_2_-scale where a higher value corresponds to higher protein expression. For each sample and assay, NPX was calculated using the following equations: 1) Ct _analyte_ – Ct _extension control_ = δCt _analyte_, 2) δCt _analyte_ – Ct _median IPC_ = δδCt _analyte_, and 3) CF _analyte_ – δδCt _analyte_ = NPX _analyte_, where Ct is the cycle threshold and CF is the correction factor. Quality control was performed in two steps: first, evaluating the standard deviation of the controls (< 0.2 for acceptance), and second, assessing individual samples’ deviations from run medians (< 0.3 NPX for acceptance).

### MPO immunofluorescence staining

To stain for the granulocyte marker myeloperoxidase (MPO), 4 µm sections of colon tissue were mounted onto Superfrost^®^ Plus Microscope slides and then baked for 3 hours at 60 °C. This was followed by sequential washes with histological-grade xylene, ethanol and double-distilled water to deparaffinize and rehydrate the tissue. A microwave was used for heat-induced epitope retrieval. Tissue slides were then boiled for 15 minutes in an alkaline Tris-based antigen-unmasking solution (Vector Laboratories, Inc., Newark, CA) containing 0.1% Tween^TM^ 20 at a pH of 9.0. Slides were subsequently rinsed in hot deionized water, transferred to a slightly acidic, citrate-based antigen-unmasking solution at a pH of 6.0 (Vector Laboratories, Inc., Newark, CA), and allowed to cool till they reached room temperature. Following antigen retrieval, slides were washed in PBS and counterstained with DAPI. Additional washes in PBS, deionized water, and Roche reaction buffer were conducted before loading the slides onto a Ventana Discovery Ultra automated slide stainer (Roche Tissue Diagnostics, Basel, Switzerland). On an autostainer, slides underwent sequential rounds of blocking, primary and secondary antibody incubations that are interspersed with washes, followed by color development. Roche protocols for denaturation and neutralization were applied between staining rounds, as required, to remove the initial antibody and quench residual horseradish peroxidase activity. After staining, the slides were again counterstained with DAPI using the autostainer. The slides were then manually washed in alternating cycles of deionized water containing 0.1% Dawn dish soap and plain deionized water, repeated for a total of five cycles. Slides were permanently mounted and allowed to dry overnight. Finally, the slides were digitally imaged at 20x magnification using a Zeiss Axio Scan.Z1 scanner (Carl Zeiss Inc., Oberkochen, Germany), and image analysis was conducted with HALO^®^ Highplex FL software (v 4.1.3; Indica Labs, Albuquerque, NM).

### H&E staining

Colonic tissue samples were collected and fixed in Z-Fix zinc formalin (Anatech Ltd., Battle Creek, MI) for 48 hours before being washed and dehydrated using a Thermo Scientific Excelsior AS^TM^ Tissue Processor (Thermo Fisher Scientific, Waltham, MA). The tissues were then transferred to a Thermo Shandon^TM^ Histocenter 3 embedding station (Thermo Fisher Scientific, Waltham, MA) where they were embedded in blocks using warm paraffin. Following this, tissue blocks were cut into 5 µm sections and baked overnight at 60 °C. This was followed by deparaffinization and rehydration through sequential exposures to xylene, ethanol, and double-distilled water. A Leica ST5010 Autostainer XL (Leica Biosystems, Nussloch, Germany) was used to complete this process. The slides were then stained with hematoxylin and counterstained with eosin Y, followed by imaging using a NanoZoomer S360 Digital Slide Scanner (Hamamatsu Photonic, Shizuoka, Japan). The images were analyzed by a board-certified veterinary pathologist using HALO^®^ Highplex FL software (v 4.1.3; Indica Labs, Albuquerque, NM).

### ELISA

Detection of sCD14 in monkey plasma was carried out using a CD14 Quantikine ELISA kit specific to rhesus macaques (R&D Systems, Minneapolis, MN) according to the manufacturer’s instructions. Plasma levels of IFN-α and IFN-γ were measured using Non- human IFN-alpha (Invitrogen, Waltham, MA) and Rhesus Macaque IFN-gamma (Novus Biologicals, CO) ELISA kits, respectively. Optical density (OD) measurements were obtained using Promega GloMax^®^ Explorer Microplate Reader (Promega Corporation, Madison, WI), and levels of CD14 and IFN-α/γ were interpolated using standard curves. For IFN-α/γ, the results were further log_2_-normalized and incorporated into the analysis presented in the O-link cytokine/chemokine volcano plot (see, **Fig. 2H**).

### Plasma metabolomics

Elucidation of plasma metabolites was done through liquid chromatography-mass spectrometry (LC-MS). For this purpose, plasma extracted prior to PBMC isolation was added to a metabolite extraction plate, which contains a sorbent filter capable of protein precipitation and removing phospholipids. This was followed by the addition of LC-MS grade pure acetonitrile (ACN) containing 0.1% (v/v) formic acid (Millipore Sigma) to the plate at an ACN-to-sample volume ratio of 3:1. The resulting mixture was then forced through the sorbent filter via the application of continuous positive pressure. This process eliminated the protein and phospholipid impurities, leaving behind samples with the necessary metabolites. The sample was then dried to a final volume of 150 µL for subsequent acquisition. Data acquisition was performed using an Orbitrap™ IQ-X™ Tribrid™ Mass Spectrometer (Thermo Fisher Scientific, Waltham, MA) fitted to a Vanquish UHPLC (ultra-high performance liquid chromatography) system (Thermo Fisher Scientific, Waltham, MA). A sample injection volume of 2 µL was used for all LC-MS runs. Samples were also subjected to reverse-phase liquid chromatography (RP-LC) through a 3 µm Discovery^®^ HS F5 HPLC column (Sigma Aldrich, St. Louis, MO) for 15 minutes. Mobile phase for these LC-MS runs consisted of 0.1% formic acid with both water (aqueous/polar) and ACN (organic/non-polar) components. The instruments were calibrated before each run to ensure data accuracy and minimize batch effects. Metabolite intensity data were quantified by calculating the area under each chromatographic peak and adjusting the values according to the sample volume.

### Statistical and data analyses

Statistical analyses were performed using GraphPad Prism (v 10.3.1; LaJolla, CA) and RStudio (2024.04.0/Build 735). Wilcoxon matched-pairs signed-rank test was used for intra-group comparisons, Mann-Whitney U-tests were used to assess differences between groups, Spearman’s non-parametric correlation was used for all correlative analyses, and PERMANOVA was used for non-parametric ANOVA with permutations. All heatmaps, PCA, and volcano plots were generated using RStudio. MetaboAnalyst 6.0 (https://www.metaboanalyst.ca/) was used for performing VIP scoring, PLS-DA, and enrichment analyses of metabolomics data, and also to generate metabolite correlation graphs. R (v 4.2.2) was used for the LASSO regression analysis of metabolomics data. A significance level of 0.05 was used consistently across all hypothesis testing. Graphical abstract and study design images were generated using BioRender.

## AUTHOR CONTRIBUTIONS

J.C.M. designed the study. J. C. M., N.S.B., C.P. (Perdios), M.H., C.D.C., and A.T.B. participated in tissue acquisition and processing. BFK provided SIVmac239M used for infection. N.S.B., C.P. (Perdios), C.D.C., and A.T.B. performed experiments. J.C.M. and N.S.B. performed the data analysis and interpretation. C. A. did the virus titer quantification. P.K., C.Z., and A.L. did plasma metabolomics. C.M.F. and B.F.K performed SIVmac239 barcode sequencing and aided with analysis. R.V.B. and A.A.S. performed histological staining. M.M. aided with flow cytometry. C.D.C., C.P. (Perdios), A.T.B., and C.P. (Palmer) provided critical and substantive intellectual editing. J.C.M. and N.S.B. wrote the manuscript. All authors approved the manuscript.

## ACKNOWLEDGMENTS

This study was generously supported by the NIH-funded TNPRC base grant P51 OD011104 and NIH grants R01 AI167644 and R21 OD031229. NIH S10 OD026800 supports the TNPRC Flow Cytometry Core. Pathogen Detection and Quantification Core (PDQC) was funded by the base grant OD011104 and the HCRPC by P51 OD011104 from the NIH. The following Research Resource Identifier (RRID):SCR-supported core facilities were utilized: Flow Cytometry Core (024611), PDQC (024614), and HCRPC (024612). Additionally, this project has been funded in whole or in part with federal funds from the National Cancer Institute, National Institutes of Health, under Contract No. 75N91019D00024. The content of this publication does not necessarily reflect the views or policies of the Department of Health and Human Services, nor does mention of trade names, commercial products, or organizations imply endorsement by the U.S. Govern-ment.

## REAGENTS RESOURCES TABLE

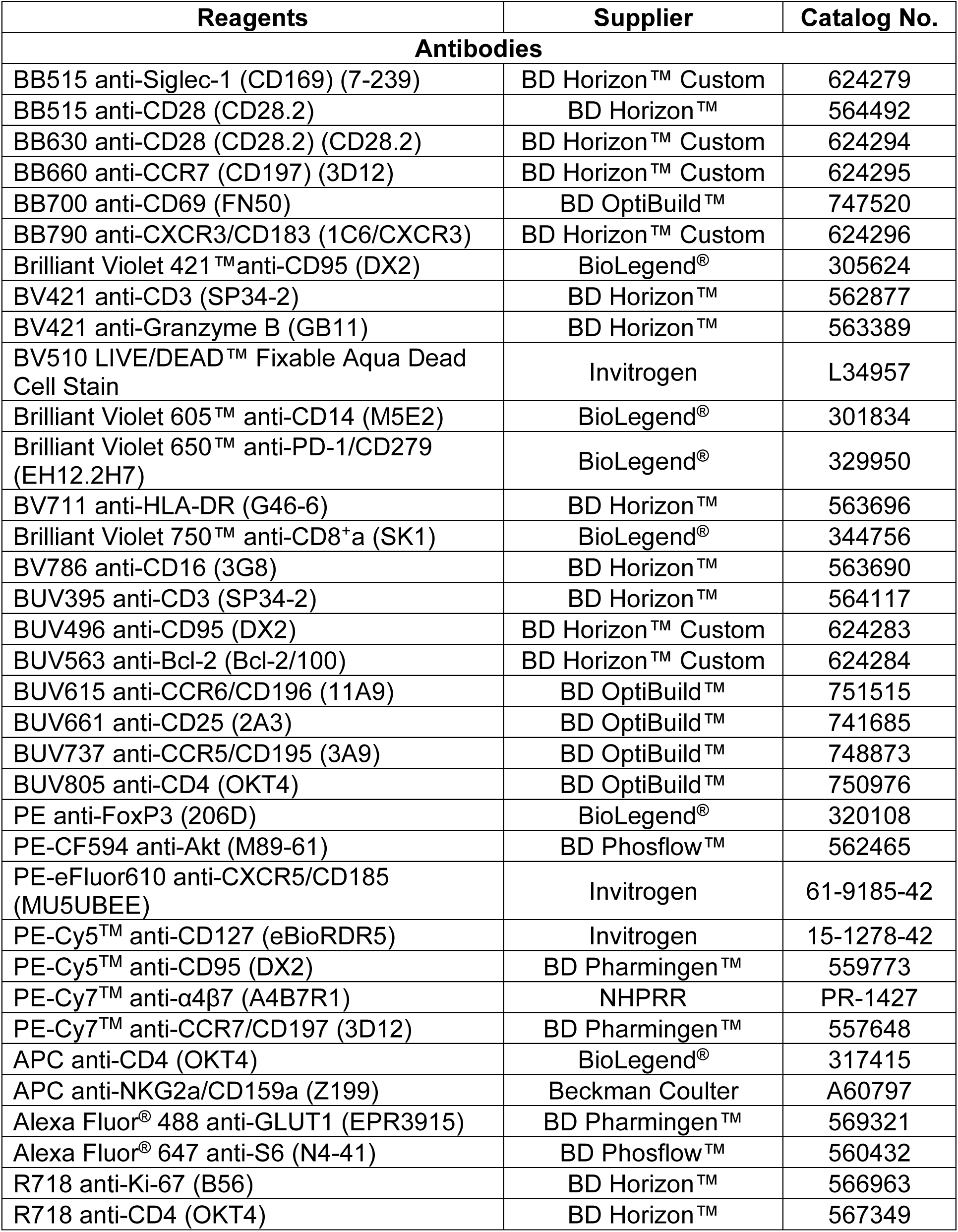
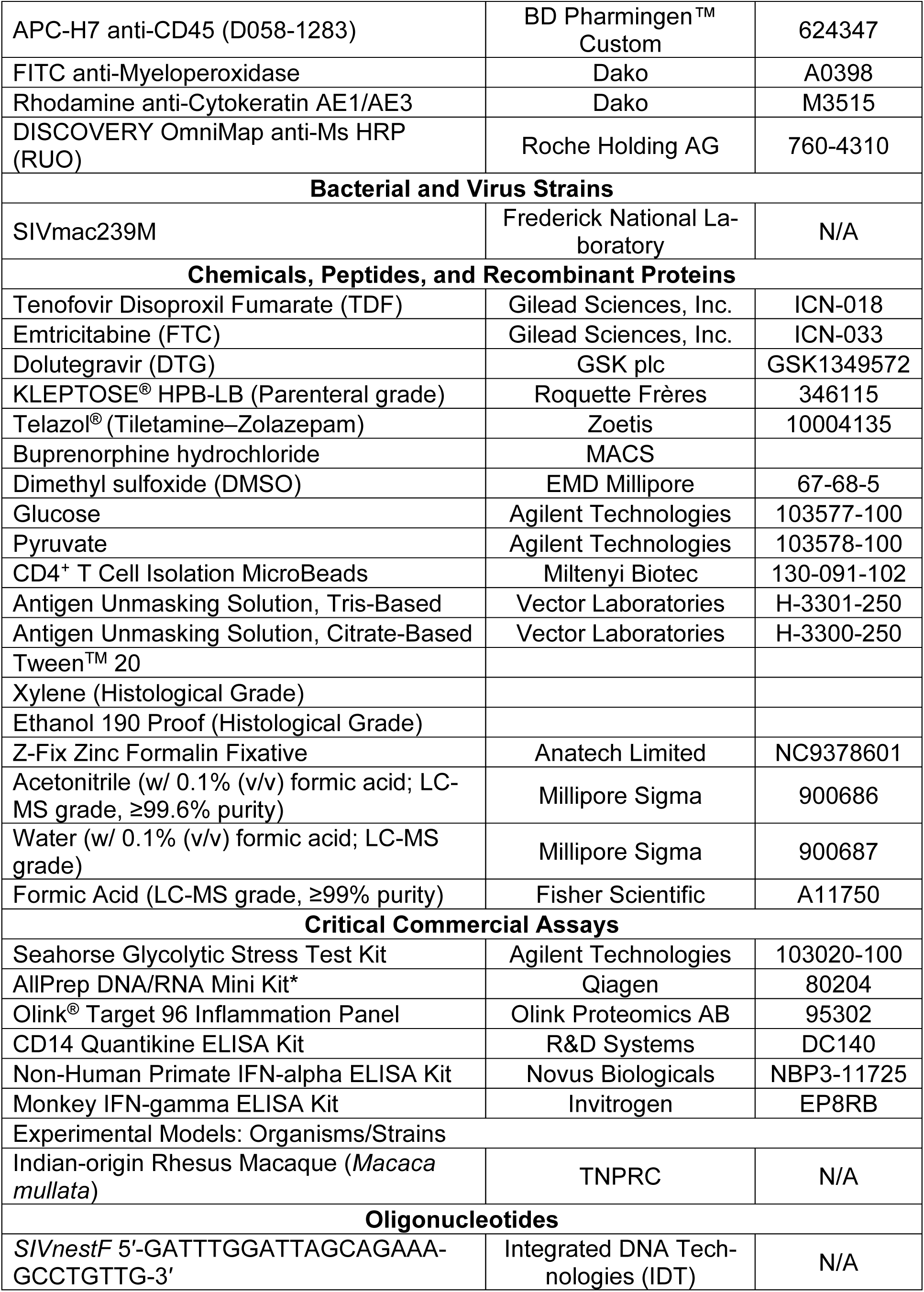
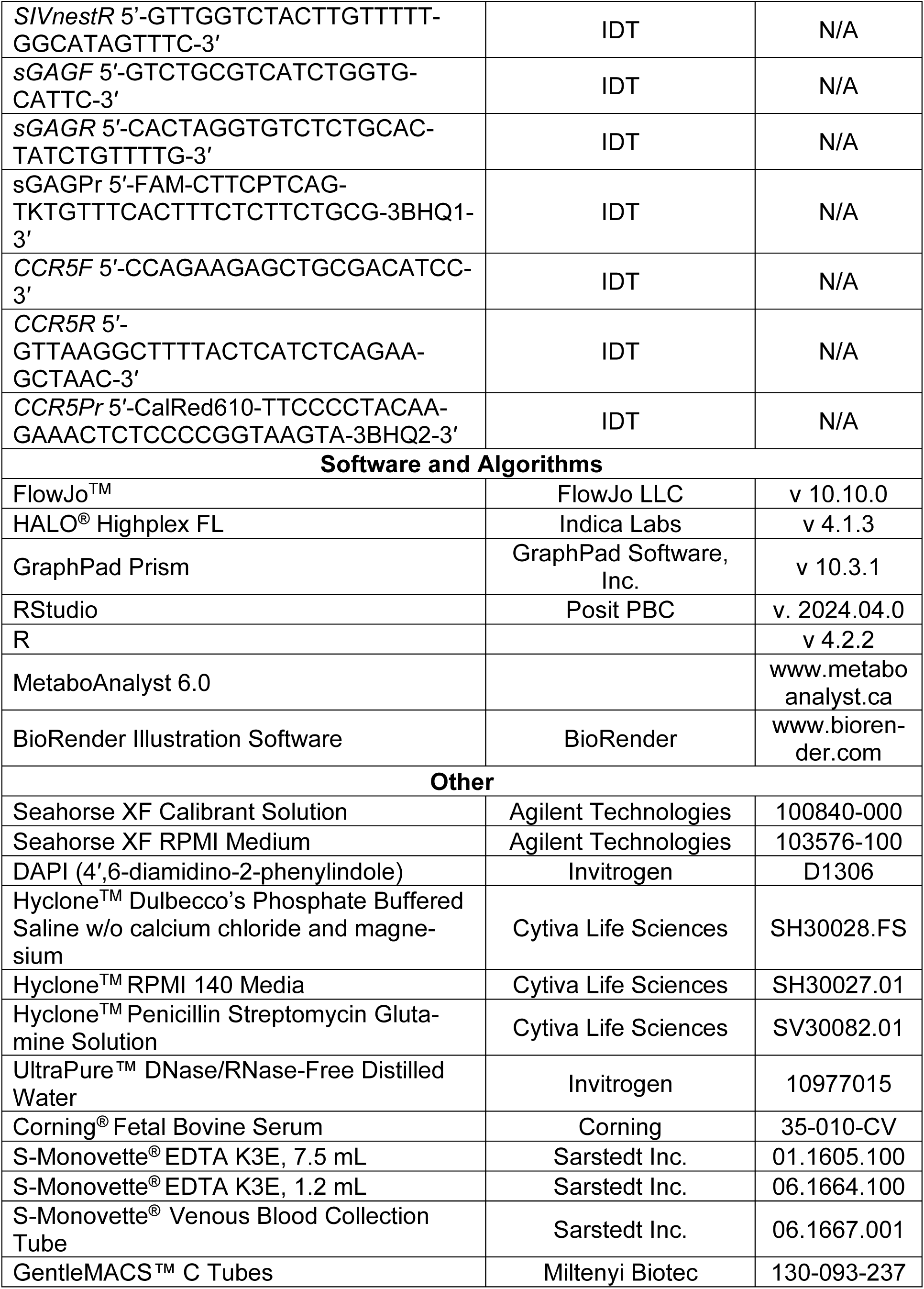
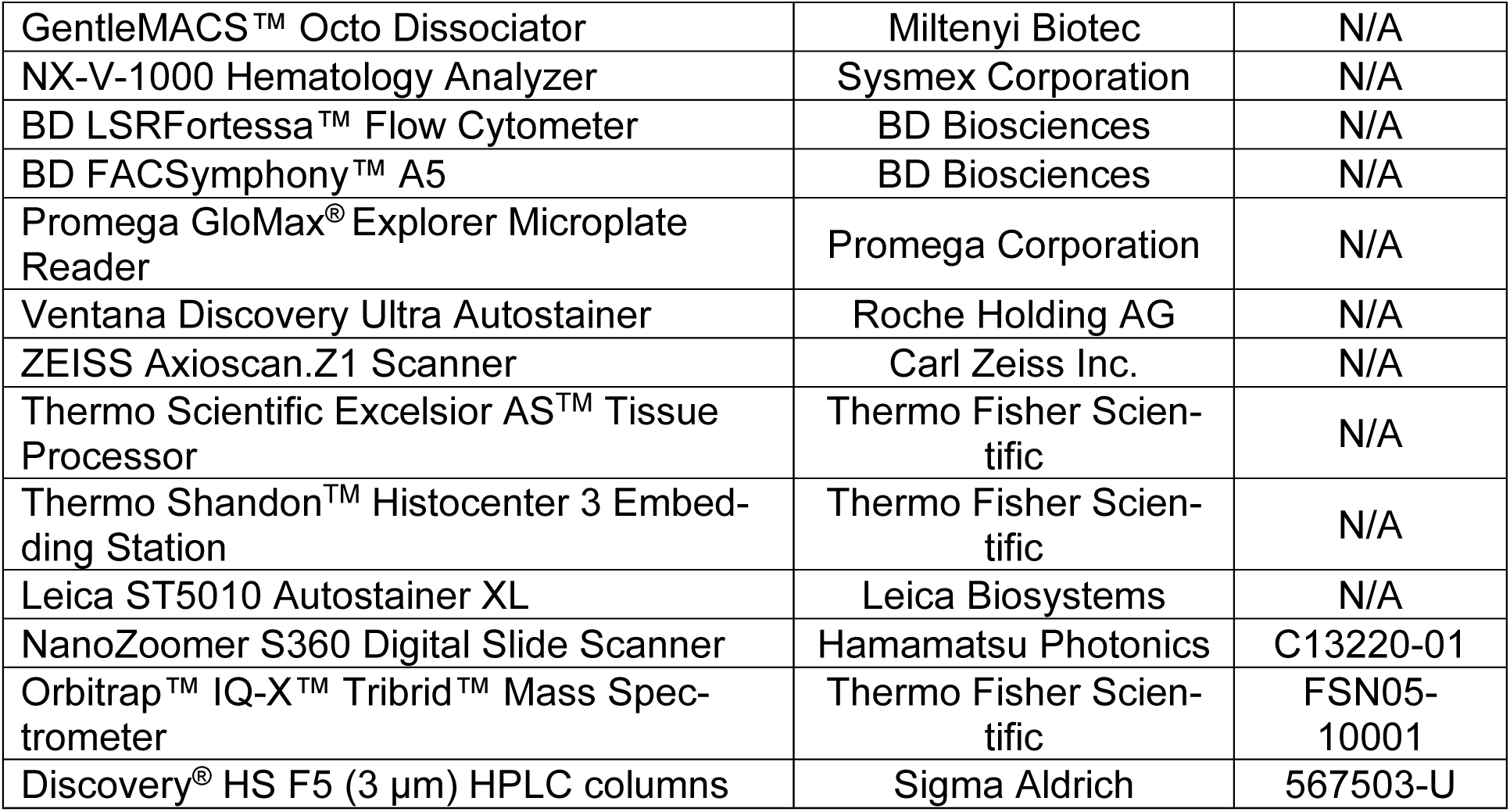

## REFERENCES

1 Chung, C. Y., Alden, S. L., Funderburg, N. T., Fu, P. & Levine, A. D. Progressive proximal-to-distal reduction in expression of the tight junction complex in colonic epithelium of virally-suppressed HIV+ individuals. PLoS Pathog 10, e1004198 (2014). 10.1371/journal.ppat.1004198

2 Brenchley, J. M. et al. CD4+ T cell depletion during all stages of HIV disease occurs predominantly in the gastrointestinal tract. J Exp Med 200, 749–759 (2004). 10.1084/jem.20040874

3 Li, Q. et al. Peak SIV replication in resting memory CD4+ T cells depletes gut lamina propria CD4+ T cells. Nature 434, 1148–1152 (2005). 10.1038/nature03513

4 Hunt, P. W. et al. Gut epithelial barrier dysfunction and innate immune activation predict mortality in treated HIV infection. J Infect Dis 210, 1228–1238 (2014). 10.1093/infdis/jiu238

5 Takata, H. et al. An active HIV reservoir during ART is associated with maintenance of HIV-specific CD8(+) T cell magnitude and short-lived differentiation status. Cell Host Microbe 31, 1494–1506 e1494 (2023). 10.1016/j.chom.2023.08.012

6 Betts, M. R. et al. HIV nonprogressors preferentially maintain highly functional HIV-specific CD8+ T cells. Blood 107, 4781–4789 (2006). 10.1182/blood-2005-12-4818

7 Champagne, P. et al. Skewed maturation of memory HIV-specific CD8 T lymphocytes. Nature 410, 106–111 (2001). 10.1038/35065118

8 McMyn, N. F. et al. The latent reservoir of inducible, infectious HIV-1 does not decrease despite decades of antiretroviral therapy. J Clin Invest 133 (2023). 10.1172/JCI171554

9 Wang, R. et al. The transcription factor Myc controls metabolic reprogramming upon T lymphocyte activation. Immunity 35, 871–882 (2011). 10.1016/j.immuni.2011.09.021

10 Bailis, W. et al. Distinct modes of mitochondrial metabolism uncouple T cell differentiation and function. Nature 571, 403–407 (2019). 10.1038/s41586-019-1311-3

11 Swamy, M. et al. Glucose and glutamine fuel protein O-GlcNAcylation to control T cell self-renewal and malignancy. Nat Immunol 17, 712–720 (2016). 10.1038/ni.3439

12 Peng, M. et al. Aerobic glycolysis promotes T helper 1 cell differentiation through an epigenetic mechanism. Science 354, 481–484 (2016). 10.1126/science.aaf6284

13 Martinez-Reyes, I. & Chandel, N. S. Mitochondrial TCA cycle metabolites control physiology and disease. Nat Commun 11, 102 (2020). 10.1038/s41467-019-13668-3

14 Chang, C. H. et al. Posttranscriptional control of T cell effector function by aerobic glycolysis. Cell 153, 1239–1251 (2013). 10.1016/j.cell.2013.05.016

15 Valle-Casuso, J. C. et al. Cellular Metabolism Is a Major Determinant of HIV-1 Reservoir Seeding in CD4(+) T Cells and Offers an Opportunity to Tackle Infection. Cell Metab 29, 611–626 e615 (2019). 10.1016/j.cmet.2018.11.015

16 Guo, H. et al. Multi-omics analyses reveal that HIV-1 alters CD4(+) T cell immunometabolism to fuel virus replication. Nat Immunol 22, 423–433 (2021). 10.1038/s41590-021-00898-1

17 Clerc, I. et al. Entry of glucose- and glutamine-derived carbons into the citric acid cycle supports early steps of HIV-1 infection in CD4 T cells. Nat Metab 1, 717–730 (2019). 10.1038/s42255-019-0084-1

18 Taylor, H. E. et al. Phospholipase D1 Couples CD4+ T Cell Activation to c-Myc-Dependent Deoxyribonucleotide Pool Expansion and HIV-1 Replication. PLoS Pathog 11, e1004864 (2015). 10.1371/journal.ppat.1004864

19 Trautmann, L. et al. Profound metabolic, functional, and cytolytic differences characterize HIV-specific CD8 T cells in primary and chronic HIV infection. Blood 120, 3466–3477 (2012). 10.1182/blood-2012-04-422550

20 Deguit, C. D. T. et al. Some Aspects of CD8+ T-Cell Exhaustion Are Associated With Altered T-Cell Mitochondrial Features and ROS Content in HIV Infection. J Acquir Immune Defic Syndr 82, 211–219 (2019). 10.1097/QAI.0000000000002121

21 Tawakol, A. et al. Association of Arterial and Lymph Node Inflammation With Distinct Inflammatory Pathways in Human Immunodeficiency Virus Infection. JAMA Cardiol 2, 163–171 (2017). 10.1001/jamacardio.2016.4728

22 Palmer, C. S. et al. Increased glucose metabolic activity is associated with CD4+ T-cell activation and depletion during chronic HIV infection. AIDS 28, 297–309 (2014). 10.1097/QAD.0000000000000128

23 Younes, S. A. et al. Cycling CD4+ T cells in HIV-infected immune nonresponders have mitochondrial dysfunction. J Clin Invest 128, 5083–5094 (2018). 10.1172/JCI120245

24 Tarancon-Diez, L. et al. Immunometabolism is a key factor for the persistent spontaneous elite control of HIV-1 infection. EBioMedicine 42, 86–96 (2019). 10.1016/j.ebiom.2019.03.004

25 Giron, L. B. et al. Non-invasive plasma glycomic and metabolic biomarkers of post-treatment control of HIV. Nat Commun 12, 3922 (2021). 10.1038/s41467-021-24077-w

26 Alrubayyi, A. et al. Functional Restoration of Exhausted CD8 T Cells in Chronic HIV-1 Infection by Targeting Mitochondrial Dysfunction. Front Immunol 13, 908697 (2022). 10.3389/fimmu.2022.908697

27 Varco-Merth, B. D. et al. Rapamycin limits CD4+ T cell proliferation in simian immunodeficiency virus-infected rhesus macaques on antiretroviral therapy. J Clin Invest 132 (2022). 10.1172/JCI156063

28 Planas, D. et al. LILAC pilot study: Effects of metformin on mTOR activation and HIV reservoir persistence during antiretroviral therapy. EBioMedicine 65, 103270 (2021). 10.1016/j.ebiom.2021.103270

29 Mansfield, K. G. et al. A diet high in saturated fat and cholesterol accelerates simian immunodeficiency virus disease progression. J Infect Dis 196, 1202–1210 (2007). 10.1086/521680

30 He, T. et al. High-fat diet exacerbates SIV pathogenesis and accelerates disease progression. J Clin Invest 129, 5474–5488 (2019). 10.1172/JCI121208

31 Mattison, J. A. et al. Caloric restriction improves health and survival of rhesus monkeys. Nat Commun 8, 14063 (2017). 10.1038/ncomms14063

32 Lakowski, B. & Hekimi, S. The genetics of caloric restriction in Caenorhabditis elegans. Proc Natl Acad Sci U S A 95, 13091–13096 (1998). 10.1073/pnas.95.22.13091

33 Palma, C. et al. Caloric Restriction Promotes Immunometabolic Reprogramming Leading to Protection from Tuberculosis. Cell Metab 33, 300–318 e312 (2021). 10.1016/j.cmet.2020.12.016

34 Mejia, P. et al. Dietary restriction protects against experimental cerebral malaria via leptin modulation and T-cell mTORC1 suppression. Nat Commun 6, 6050 (2015). 10.1038/ncomms7050

35 Collins, N. et al. The Bone Marrow Protects and Optimizes Immunological Memory during Dietary Restriction. Cell 178, 1088–1101 e1015 (2019). 10.1016/j.cell.2019.07.049

36 Chiou, K. L. et al. Rhesus macaques as a tractable physiological model of human ageing. Philos Trans R Soc Lond B Biol Sci 375, 20190612 (2020). 10.1098/rstb.2019.0612

37 Duregon, E. et al. Prolonged fasting times reap greater geroprotective effects when combined with caloric restriction in adult female mice. Cell Metab 35, 1179–1194 e1175 (2023). 10.1016/j.cmet.2023.05.003

38 Velingkaar, N. et al. Reduced caloric intake and periodic fasting independently contribute to metabolic effects of caloric restriction. Aging Cell 19, e13138 (2020). 10.1111/acel.13138

39 Berman, C. M. & Schwartz, S. A nonintrusive method for determining relative body fat in free-ranging monkeys. Am J Primatol 14, 53–64 (1988). 10.1002/ajp.1350140105

40 Tomiyama, A. J. et al. Low calorie dieting increases cortisol. Psychosom Med 72, 357–364 (2010). 10.1097/PSY.0b013e3181d9523c

41 Fontana, L. et al. Effects of 2-year calorie restriction on circulating levels of IGF-1, IGF-binding proteins and cortisol in nonobese men and women: a randomized clinical trial. Aging Cell 15, 22–27 (2016). 10.1111/acel.12400

42 Nakamura, Y., Walker, B. R. & Ikuta, T. Systematic review and meta-analysis reveals acutely elevated plasma cortisol following fasting but not less severe calorie restriction. Stress 19, 151–157 (2016). 10.3109/10253890.2015.1121984

43 Vamvini, M. T., Aronis, K. N., Chamberland, J. P. & Mantzoros, C. S. Energy deprivation alters in a leptin- and cortisol-independent manner circulating levels of activin A and follistatin but not myostatin in healthy males. J Clin Endocrinol Metab 96, 3416–3423 (2011). 10.1210/jc.2011-1665

44 Immonen, T. T. et al. Genetically barcoded SIV reveals the emergence of escape mutations in multiple viral lineages during immune escape. Proc Natl Acad Sci U S A 117, 494–502 (2020). 10.1073/pnas.1914967117

45 Veazey, R. S. et al. Gastrointestinal tract as a major site of CD4+ T cell depletion and viral replication in SIV infection. Science 280, 427–431 (1998). 10.1126/science.280.5362.427

46 Mehandru, S. et al. Primary HIV-1 infection is associated with preferential depletion of CD4+ T lymphocytes from effector sites in the gastrointestinal tract. J Exp Med 200, 761–770 (2004). 10.1084/jem.20041196

47 Raehtz, K. D. et al. African green monkeys avoid SIV disease progression by preventing intestinal dysfunction and maintaining mucosal barrier integrity. PLoS Pathog 16, e1008333 (2020). 10.1371/journal.ppat.1008333

48 Jacquelin, B. et al. Nonpathogenic SIV infection of African green monkeys induces a strong but rapidly controlled type-I IFN response. J Clin Invest 119, 3544–3555 (2009). 10.1172/JCI40093

49 Bourgoin, P., Biechele, G., Ait Belkacem, I., Morange, P. E. & Malergue, F. Role of the interferons in CD64 and CD169 expressions in whole blood: Relevance in the balance between viral- or bacterial-oriented immune responses. Immun Inflamm Dis 8, 106–123 (2020). 10.1002/iid3.289

50 Akiyama, H. et al. Interferon-Inducible CD169/Siglec1 Attenuates Anti-HIV-1 Effects of Alpha Interferon. J Virol 91 (2017). 10.1128/JVI.00972-17

51 Kim, H. J. et al. Blood monocyte-derived CD169(+) macrophages contribute to antitumor immunity against glioblastoma. Nat Commun 13, 6211 (2022). 10.1038/s41467-022-34001-5

52 Perdios, C. et al. RhCMV expands CCR5+ memory T cells and promotes SIV reservoir seeding in the gut mucosa. JCI Insight 11 (2026). 10.1172/jci.insight.198743

53 Li, G. et al. Depletion of plasmacytoid dendritic cells rescues HIV-reactive stem-like CD8(+) T cells during chronic HIV-1 infection. Sci Transl Med 17, eadr3930 (2025). 10.1126/scitranslmed.adr3930

54 Chin, Y. E. et al. Cell growth arrest and induction of cyclin-dependent kinase inhibitor p21 WAF1/CIP1 mediated by STAT1. Science 272, 719–722 (1996). 10.1126/science.272.5262.719

55 Lukhele, S. et al. The transcription factor IRF2 drives interferon-mediated CD8(+) T cell exhaustion to restrict anti-tumor immunity. Immunity 55, 2369–2385 e2310 (2022). 10.1016/j.immuni.2022.10.020

56 Sandler, N. G. et al. Type-I interferon responses in rhesus macaques prevent SIV infection and slow disease progression. Nature 511, 601–605 (2014). 10.1038/nature13554

57 Yang, H. & Yang, L. Targeting cAMP/PKA pathway for glycemic control and type 2 diabetes therapy. J Mol Endocrinol 57, R93–R108 (2016). 10.1530/JME-15-0316

58 Frey, P. A. The Leloir pathway: a mechanistic imperative for three enzymes to change the stereochemical configuration of a single carbon in galactose. FASEB J 10, 461–470 (1996).

59 van der Windt, G. J. et al. Mitochondrial respiratory capacity is a critical regulator of CD8+ T cell memory development. Immunity 36, 68–78 (2012). 10.1016/j.immuni.2011.12.007

60 Pollizzi, K. N. et al. Asymmetric inheritance of mTORC1 kinase activity during division dictates CD8(+) T cell differentiation. Nat Immunol 17, 704–711 (2016). 10.1038/ni.3438

61 Pollizzi, K. N. et al. mTORC1 and mTORC2 selectively regulate CD8(+) T cell differentiation. J Clin Invest 125, 2090–2108 (2015). 10.1172/JCI77746

62 Buller, C. L., Heilig, C. W. & Brosius, F. C., 3rd. GLUT1 enhances mTOR activity independently of TSC2 and AMPK. Am J Physiol Renal Physiol 301, F588–596 (2011). 10.1152/ajprenal.00472.2010

63 Jordan, S. et al. Dietary Intake Regulates the Circulating Inflammatory Monocyte Pool. Cell 178, 1102–1114 e1117 (2019). 10.1016/j.cell.2019.07.050

64 Sandler, N. G. et al. Plasma levels of soluble CD14 independently predict mortality in HIV infection. J Infect Dis 203, 780–790 (2011). 10.1093/infdis/jiq118

65 De Clercq, J. et al. Longitudinal patterns of inflammatory mediators after acute HIV infection correlate to intact and total reservoir. Front Immunol 14, 1337316 (2023). 10.3389/fimmu.2023.1337316

66 Dwivedi, A. K. et al. A cohort-based study of host gene expression: tumor suppressor and innate immune/inflammatory pathways associated with the HIV reservoir size. PLoS Pathog 19, e1011114 (2023). 10.1371/journal.ppat.1011114

67 Angin, M. et al. Metabolic plasticity of HIV-specific CD8(+) T cells is associated with enhanced antiviral potential and natural control of HIV-1 infection. Nat Metab 1, 704–716 (2019). 10.1038/s42255-019-0081-4

68 Pearce, E. L., Poffenberger, M. C., Chang, C. H. & Jones, R. G. Fueling immunity: insights into metabolism and lymphocyte function. Science 342, 1242454 (2013). 10.1126/science.1242454

69 Gerriets, V. A. et al. Metabolic programming and PDHK1 control CD4+ T cell subsets and inflammation. J Clin Invest 125, 194–207 (2015). 10.1172/JCI76012

70 Butterfield, T. R. et al. Increased glucose transporter-1 expression on intermediate monocytes from HIV-infected women with subclinical cardiovascular disease. AIDS 31, 199–205 (2017). 10.1097/QAD.0000000000001320

71 Brust, D. et al. Fluorodeoxyglucose imaging in healthy subjects with HIV infection: impact of disease stage and therapy on pattern of nodal activation. AIDS 20, 495–503 (2006). 10.1097/01.aids.0000210603.40267.29

72 Zaidi, A. & Sharma, S. Arrhythmogenic right ventricular remodelling in endurance athletes: Pandora’s box or Achilles’ heel? Eur Heart J 36, 1955–1957 (2015). 10.1093/eurheartj/ehv199

73 Wu, H. et al. T-cells produce acidic niches in lymph nodes to suppress their own effector functions. Nat Commun 11, 4113 (2020). 10.1038/s41467-020-17756-7

74 Stryhn, A. et al. pH dependence of MHC class I-restricted peptide presentation. J Immunol 156, 4191–4197 (1996).

75 Pucino, V. et al. Lactate Buildup at the Site of Chronic Inflammation Promotes Disease by Inducing CD4(+) T Cell Metabolic Rewiring. Cell Metab 30, 1055–1074 e1058 (2019). 10.1016/j.cmet.2019.10.004

76 Feng, J. et al. Tumor cell-derived lactate induces TAZ-dependent upregulation of PD-L1 through GPR81 in human lung cancer cells. Oncogene 36, 5829–5839 (2017). 10.1038/onc.2017.188

77 Hickey, J. W. et al. Organization of the human intestine at single-cell resolution. Nature 619, 572–584 (2023). 10.1038/s41586-023-05915-x

78 Guerbette, T., Boudry, G. & Lan, A. Mitochondrial function in intestinal epithelium homeostasis and modulation in diet-induced obesity. Mol Metab 63, 101546 (2022). 10.1016/j.molmet.2022.101546

79 Simonetti, F. R. et al. Antigen-driven clonal selection shapes the persistence of HIV-1-infected CD4+ T cells in vivo. J Clin Invest 131 (2021). 10.1172/JCI145254

80 Reeves, D. B. et al. A majority of HIV persistence during antiretroviral therapy is due to infected cell proliferation. Nat Commun 9, 4811 (2018). 10.1038/s41467-018-06843-5

81 Reeves, D. B. et al. Estimating the contribution of CD4 T cell subset proliferation and differentiation to HIV persistence. Nat Commun 14, 6145 (2023). 10.1038/s41467-023-41521-1

82 Arandjelovic, P. et al. Venetoclax, alone and in combination with the BH3 mimetic S63845, depletes HIV-1 latently infected cells and delays rebound in humanized mice. Cell Rep Med 4, 101178 (2023). 10.1016/j.xcrm.2023.101178

83 Collora, J. A. et al. Single-cell multiomics reveals persistence of HIV-1 in expanded cytotoxic T cell clones. Immunity 55, 1013–1031 e1017 (2022). 10.1016/j.immuni.2022.03.004

84 Pape, K. A., Khoruts, A., Mondino, A. & Jenkins, M. K. Inflammatory cytokines enhance the in vivo clonal expansion and differentiation of antigen-activated CD4+ T cells. J Immunol 159, 591–598 (1997).

85 Del Prete, G. Q. et al. TLR7 agonist administration to SIV-infected macaques receiving early initiated cART does not induce plasma viremia. JCI Insight 4 (2019). 10.1172/jci.insight.127717

86 Dashti, A. et al. AZD5582 plus SIV-specific antibodies reduce lymph node viral reservoirs in antiretroviral therapy-suppressed macaques. Nat Med 29, 2535–2546 (2023). 10.1038/s41591-023-02570-7

87 Di Francesco, A. et al. Dietary restriction impacts health and lifespan of genetically diverse mice. Nature 634, 684–692 (2024). 10.1038/s41586-024-08026-3

88 Shah, M. & Vella, A. Effects of GLP-1 on appetite and weight. Rev Endocr Metab Disord 15, 181–187 (2014). 10.1007/s11154-014-9289-5

89 Khanal, S. et al. In Vivo Validation of the Viral Barcoding of Simian Immunodeficiency Virus SIVmac239 and the Development of New Barcoded SIV and Subtype B and C Simian-Human Immunodeficiency Viruses. J Virol 94 (2019). 10.1128/JVI.01420-19

90 Pruessner, J. C., Kirschbaum, C., Meinlschmid, G. & Hellhammer, D. H. Two formulas for computation of the area under the curve represent measures of total hormone concentration versus time-dependent change. Psychoneuroendocrinology 28, 916–931 (2003). 10.1016/s0306-4530(02)00108-7

91 Monjure, C. J. et al. Optimization of PCR for quantification of simian immunodeficiency virus genomic RNA in plasma of rhesus macaques (Macaca mulatta) using armored RNA. J Med Primatol 43, 31–43 (2014). 10.1111/jmp.12088

92 Hansen, S. G. et al. Profound early control of highly pathogenic SIV by an effector memory T-cell vaccine. Nature 473, 523–527 (2011). 10.1038/nature10003

93 Fennessey, C. M. et al. Genetically-barcoded SIV facilitates enumeration of rebound variants and estimation of reactivation rates in nonhuman primates following interruption of suppressive antiretroviral therapy. PLoS Pathog 13, e1006359 (2017). 10.1371/journal.ppat.1006359

94 Assarsson, E. et al. Homogenous 96-plex PEA immunoassay exhibiting high sensitivity, specificity, and excellent scalability. PLoS One 9, e95192 (2014). 10.1371/journal.pone.0095192

