## Supplemental Figures & Tables for "Metabolic reprogramming by caloric restriction enhances acute phase virological control and reduces chronic inflammation in SIV-infected rhesus macaques"

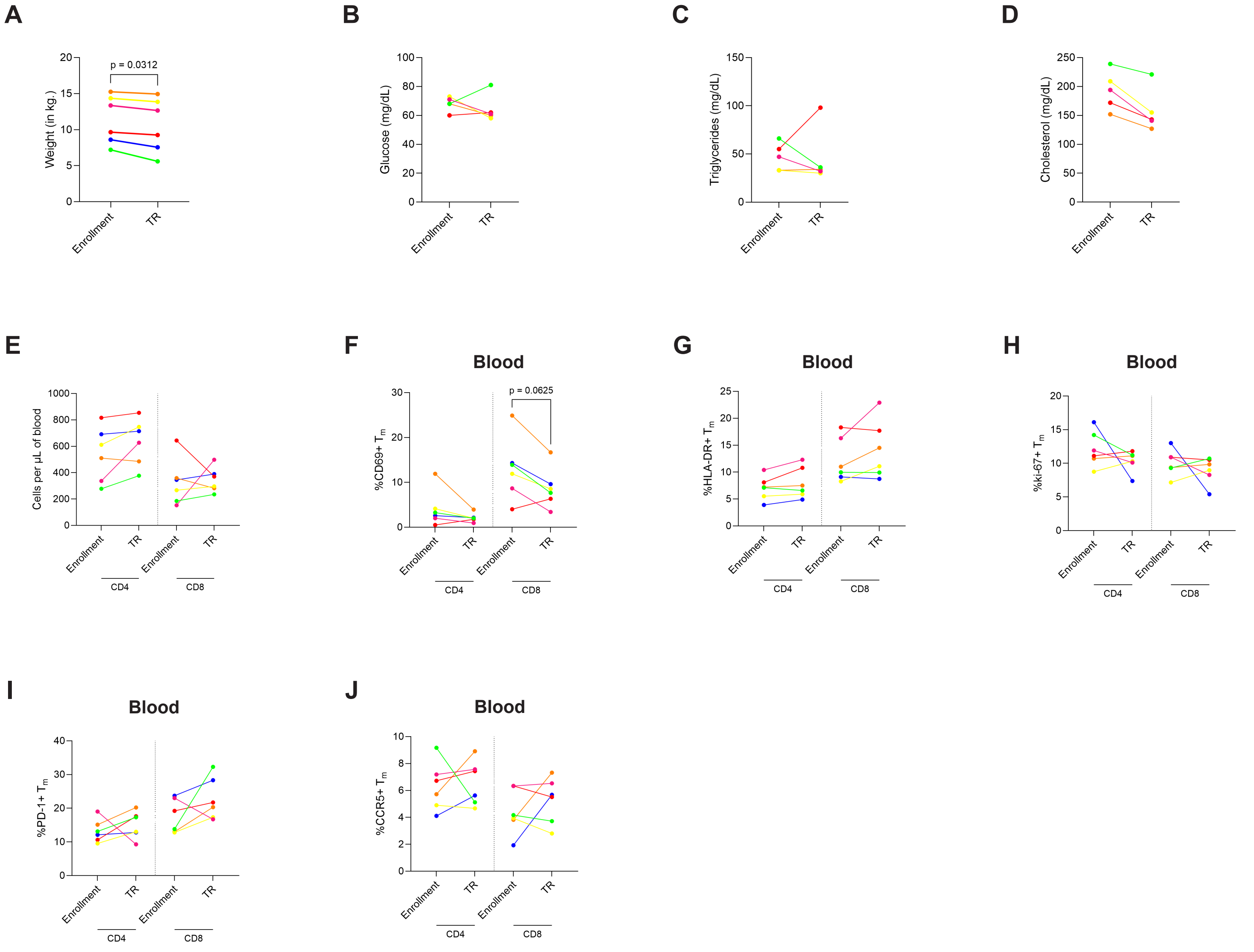


**Supplementary Figure 1. Time restricted (TR) feeding regimen modulates activation and proliferation marker expression in memory T cells.**

**(A)** Weight (in kg.) at study enrollment and following TR.

**(B-D)** Plasma levels (mg/dL) of glucose **(B)**, triglycerides **(C)**, and cholesterol **(D)** at study enrollment and following TR. RM01 is not included at TR time point as serum was not collected for blood chemistry panel.

**(E)** Absolute counts (cells/µL) of total $CD4^{+}$ and $CD8^{+}$ T cells in the blood at enrollment and following TR. Counts were calculated using total lymphocyte counts from complete blood chemistries and multiplied by frequencies of viable $CD3^{+}$ $CD4^{+}$ or $CD8^{+}$ lymphocytes assessed via flow cytometry.

**(F-J)** Percentage of $CD4^{+}$ and $CD8^{+}$ memory T cells expressing CD69 **(F)**, HLA-DR **(G)**, Ki-67 **(H)**, PD-1 **(I)**, and CCR5 **(J)** at study enrollment and TR.

**Statistical analyses: (A-J):** Wilcoxon matched-pairs signed rank test. α=0.05.

**A**

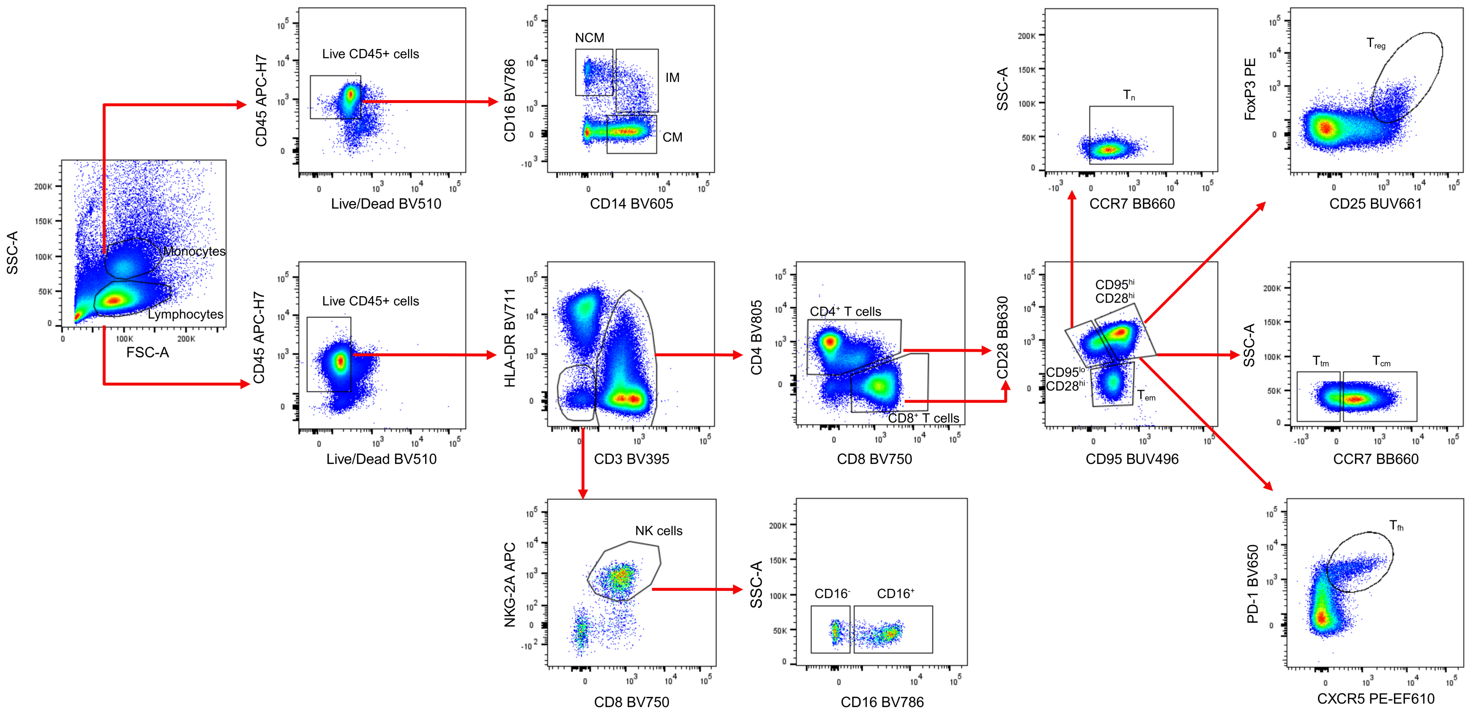


**Supplementary Figure 2. Gating strategy used to define key immune cell subsets through a 28-color flow cytometry panel.**

**(A)** Distinct immune cell subsets were defined are as follows: naïve ($T_n$)($CD28^+$ $CD95^-$ $CCR7^+$); central memory ($T_{cm}$)($CD28^+$ $CD95^+$ $CCR7^+$); transitional memory ($T_{tm}$)($CD28^+$ $CD95^+$ $CCR7^-$); effector memory ($T_{em}$)($CD28^-$ $CD95^+$ $CCR7^-$); $CD4^+$ T regulatory ($T_{reg}$)($CD28^+$ $CD95^+$ $CD25^+$ $FoxP3^+$); $CD4^+$ T follicular helper (pLN only)($T_{fh}$)($CD28^+$ $CD95^+$ $CXCR5^+$ $PD\text{-}1^+$); Natural Killer (NK) cell ($CD3^-$ $HLA\text{-}DR^-$ $NKG2A^+$ $CD8a^+$ $CD16^{+/-}$); CM, classical monocyte ($CD14^+$ CD16-); IM, intermediate monocyte ($CD14^+$ $CD16^+$); NCM, non-classical monocyte ($CD14^{dim}$ $CD16^+$). The depicted dot plots are representative of a PBMC sample.

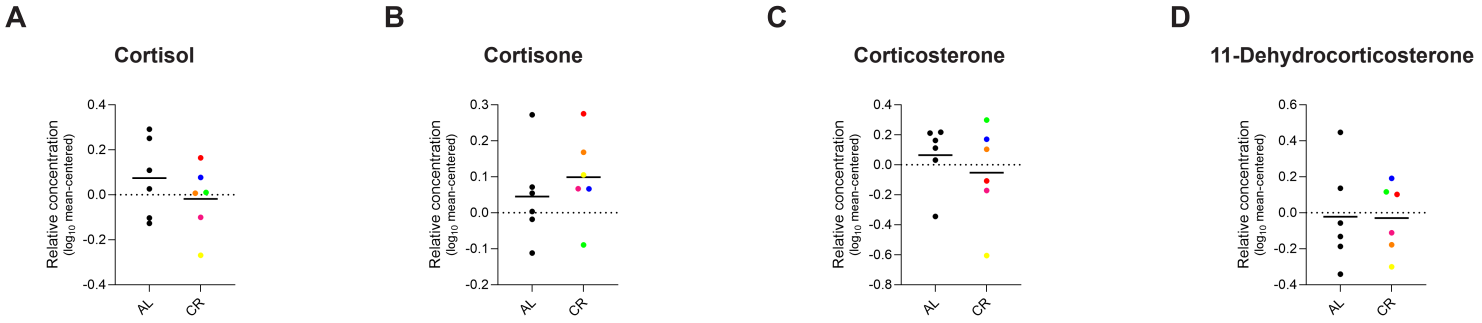


**Supplementary Figure 3. CR does not elevate plasma concentrations of glucocorticoid stress hormones prior to SIV infection.**

**(A-D)** Relative plasma concentrations (mean-centered $\log_{10}$ normalized) of cortisol **(A),** cortisone **(B)**, corticosterone **(C)** and 11-Dehydrocorticosterone **(D)** in AL and CR animals prior to SIV infection, as measured by LC-MS metabolomics.

**Statistical analyses: (A-D):** Mann-Whitney U test. α=0.05.

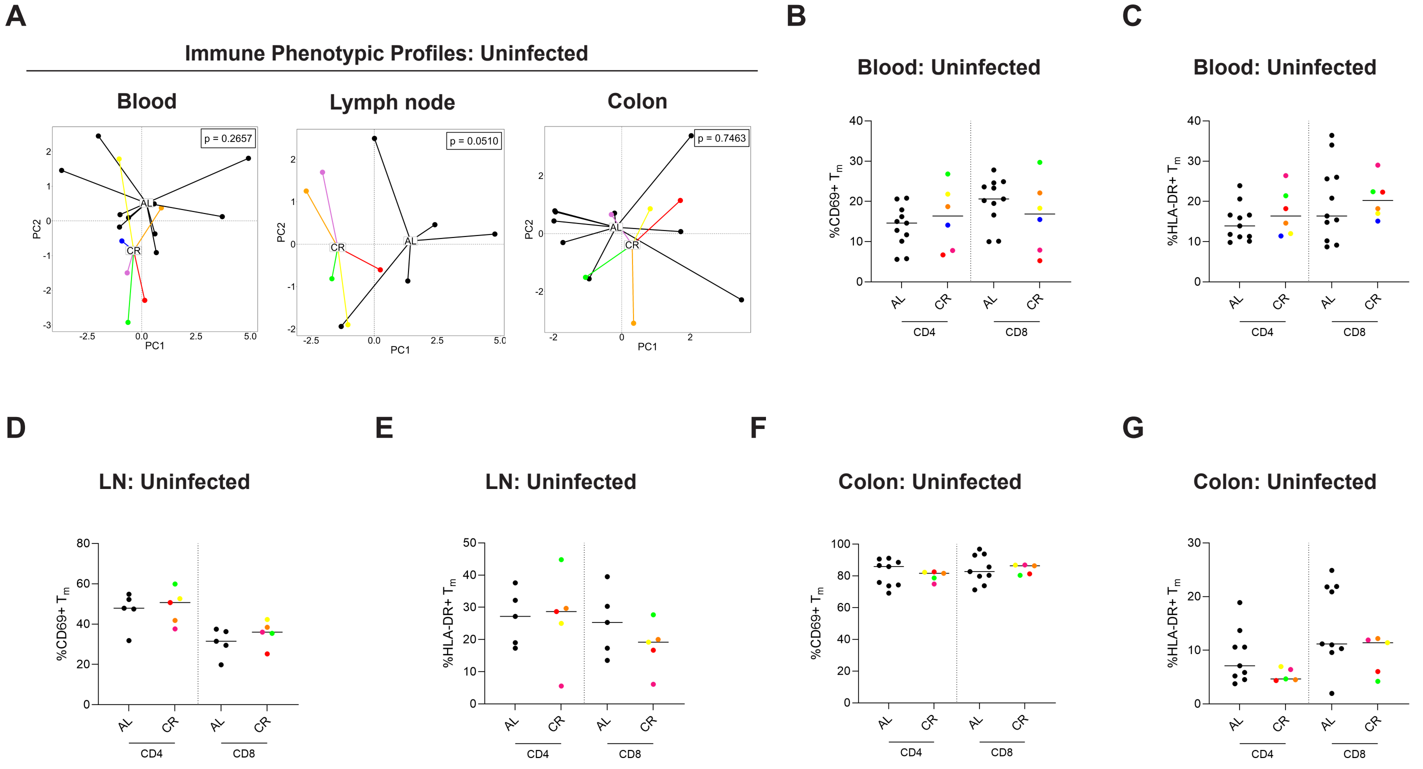


**Supplementary Figure 4. CR alters CCR5 expression on peripheral CD4$^+$ and CD8$^+$ T cells without broadly impacting the overall immune phenotypic profile.**

**(A)** Principal component analyses (PCA) of 15 immune phenotypic markers on distinct immune populations described in Supplementary figure 2 across blood, lymph nodes, and colon in AL and CR animals prior to SIV infection.

**(B & C)** Percentage of CD4$^+$ and CD8$^+$ memory T cells in the blood expressing CD69 **(B)** and HLA-DR **(C)** in AL and CR animals prior to SIV infection

**(D & E)** Percentage of CD4$^+$ and CD8$^+$ memory T cells in peripheral lymph nodes expressing CD69 **(D)** and HLA-DR **(E)** in AL and CR animals prior to SIV infection

**(F & G)** Percentage of CD4$^+$ and CD8$^+$ memory T cells in the colon expressing CD69 **(F)** and HLA-DR **(G)** in AL and CR animals prior to SIV infection.

**Statistical analyses: (A)** PERMANOVA, and **(B-G)** Mann-Whitney U test. α=0.05.

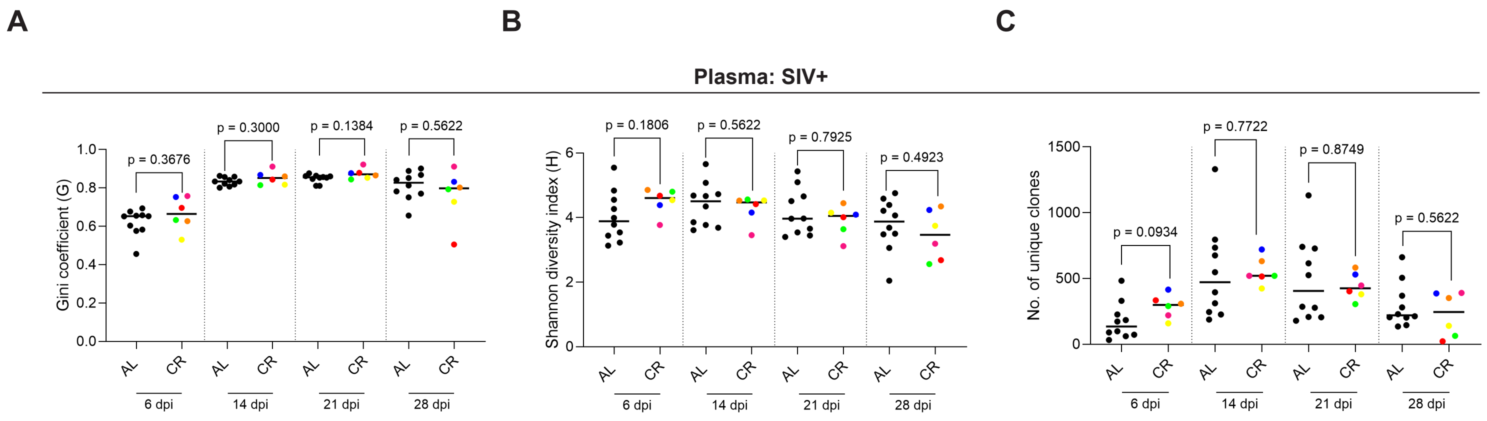


**Supplementary Figure 5. CR does not alter the diversity of early SIV viral population.**

**(A-C)** SIVmac239 barcode sequencing analysis depicting viral population diversity in the plasma of AL and CR animals at 6,14, 21 and 28 dpi. Viral diversity is represented using three indices: Gini Coefficient **(A)**, Shannon diversity index **(B)** and no. of unique viral clones **(C).**

**Statistical analyses: (A-C)** Mann-Whitney U test. α=0.05.

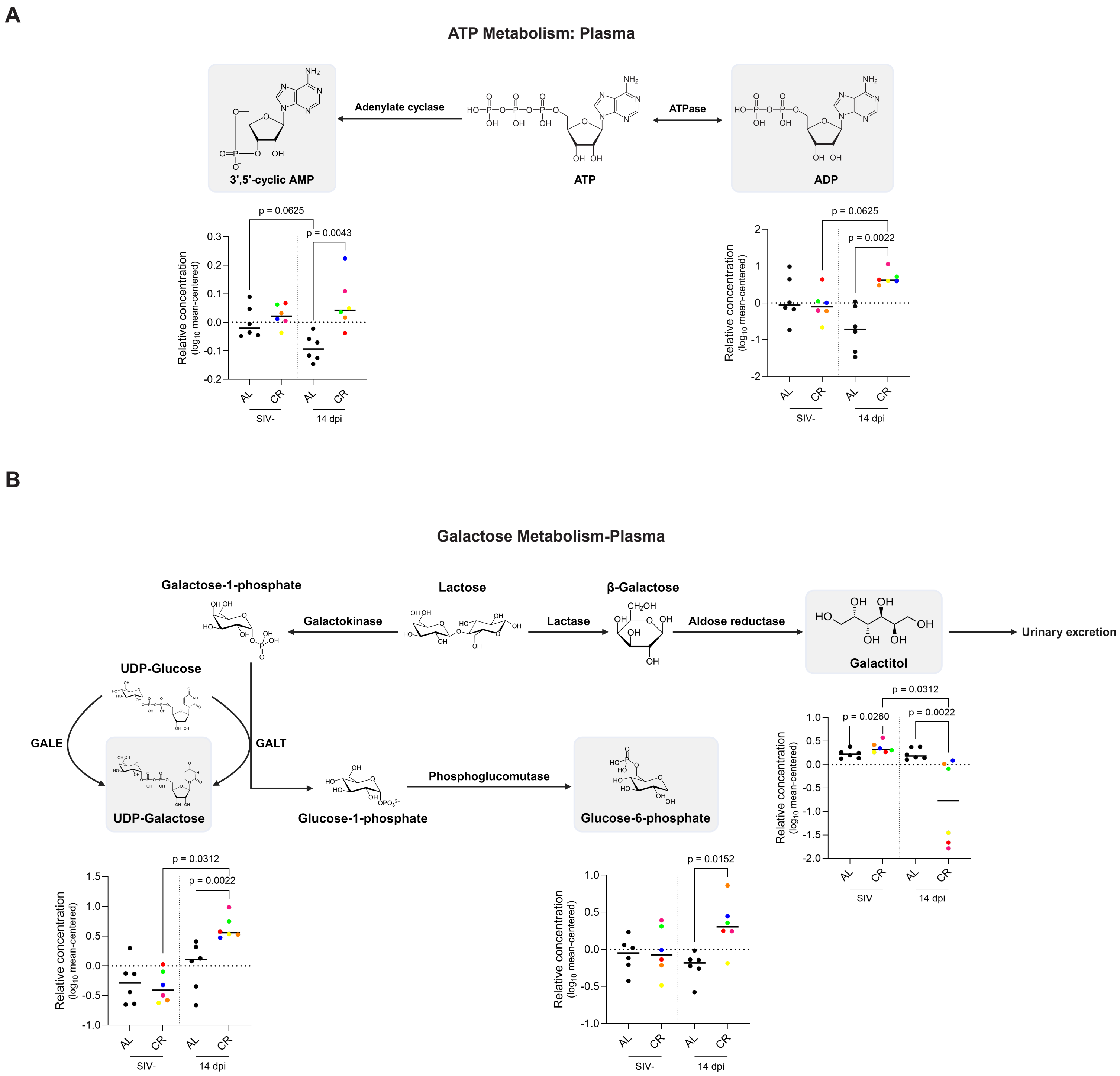


**Supplementary Figure 6. CR is associated with elevated levels of metabolites associated with ATP and galactose metabolism in the plasma following SIV infection.**

**(A)** Relative plasma concentrations of ADP and cAMP (cyclic adenosine monophosphate) in AL and CR animals at uninfected and 14 dpi time points.

**(B)** Relative plasma concentrations (mean-centered $log_{10}$ normalized) of UDP-galactose, galactitol, and G6P in AL and CR animals at uninfected and 14 dpi time points.

**Statistical analyses: (A & B)** Mann-Whitney U test and Wilcoxon signed rank test for inter- and intra-cohort comparisons, respectively. α=0.05.

B

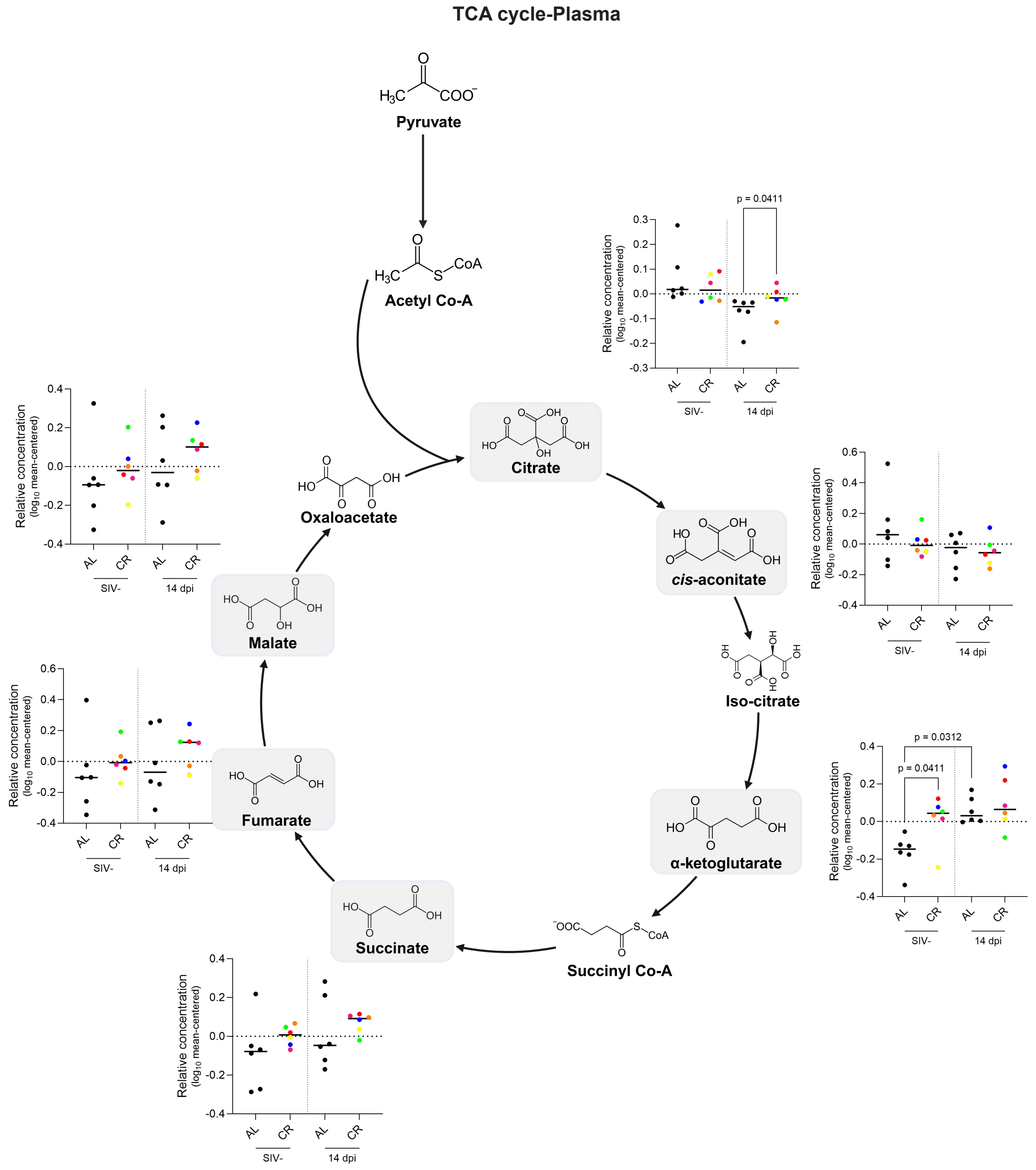
TCA cycle-Plasma
Pyruvate
Acetyl Co-A
Citrate
cis-aconitate
Iso-citrate
α-ketoglutarate
Succinyl Co-A
Succinate
Fumarate
Malate
Oxaloacetate
Relative concentration (log10 mean-centered)
p = 0.0411
p = 0.0312
AL
CR
SIV-
14 dpi

C

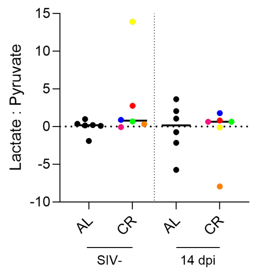
Lactate : Pyruvate
AL
CR
SIV-
14 dpi

D

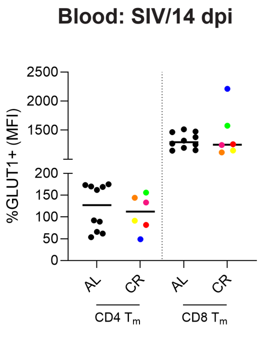
Blood: SIV/14 dpi
%GLUT1+ (MFI)
AL
CR
CD4 Tm
CD8 Tm

**Supplementary Figure 7. Plasma glycolytic profile in CR animals is independent of changes to TCA cycle or intracellular glucose flux.**

**(A)** Relative plasma concentrations of key TCA cycle intermediates in AL and CR animals before SIV infection and 14 dpi.

**(B)** Ratio of plasma lactate and pyruvate in AL and CR animals before SIV infection and 14 dpi.

**(C)** Median Fluorescence Intensity (MFI) of GLUT1 expression on $CD4^+$ and $CD8^+$ memory T cells in the blood of AL and CR animals at 14 dpi (acquired via flow cytometry).

**Statistical analyses: (A-C):** Mann-Whitney U test and Wilcoxon matched-pairs signed rank test were used for inter- and intra-cohort comparisons, respectively. $\alpha=0.05$.

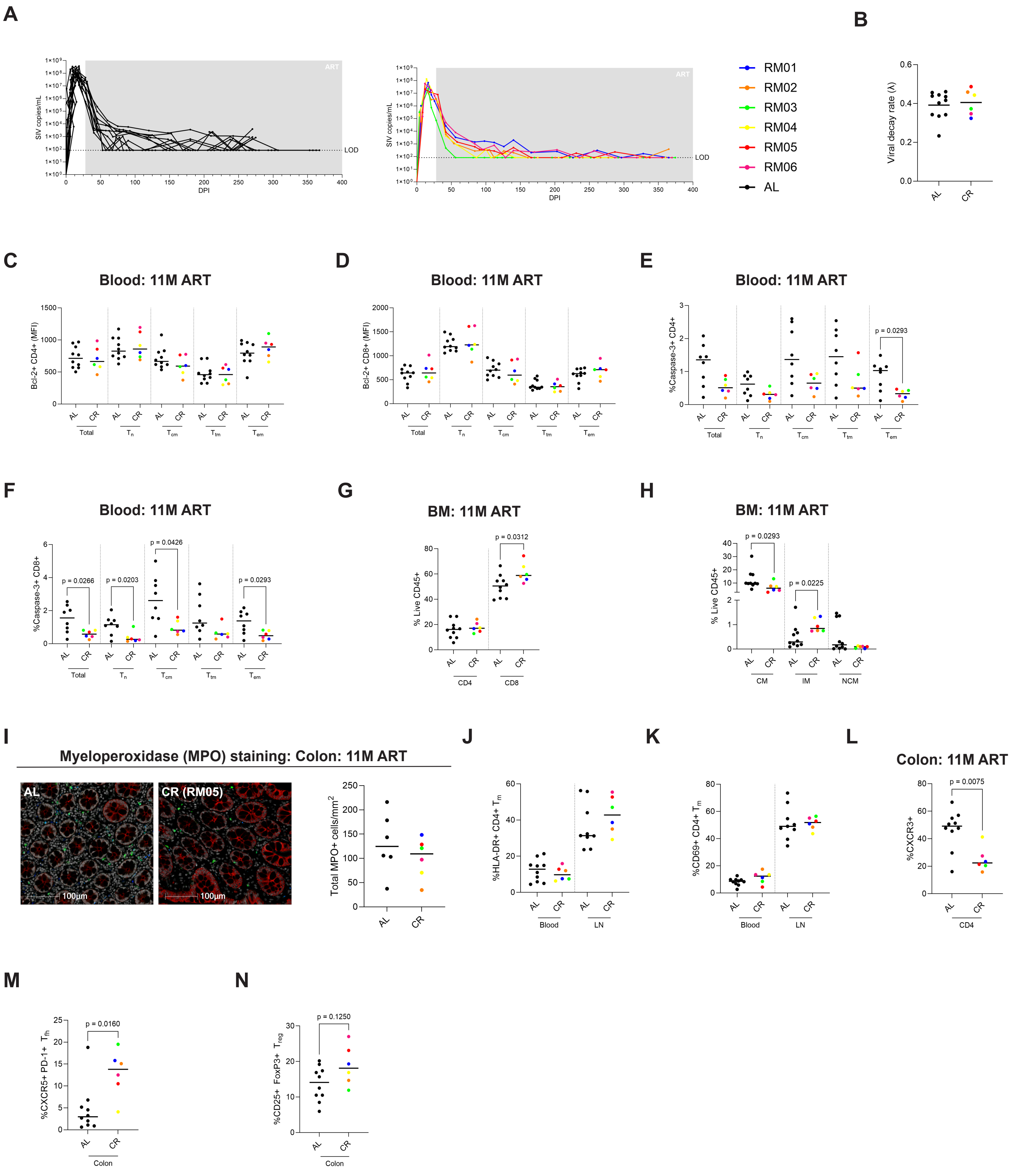
A
B
SIV copies/mL
DPI
ART
LOD
RM01
RM02
RM03
RM04
RM05
RM06
AL
Viral decay rate (λ)
AL
CR
C
D
E
F
Blood: 11M ART
Bcl-2+ CD4+ (MFI)
Bcl-2+ CD8+ (MFI)
%Caspase-3+ CD4+
%Caspase-3+ CD8+
Total
$T_n$
$T_{cm}$
$T_{tm}$
$T_{em}$
p = 0.0293
p = 0.0266
p = 0.0203
p = 0.0426
p = 0.0293
G
BM: 11M ART
% Live CD45+
CD4
CD8
p = 0.0312
H
BM: 11M ART
% Live CD45+
CM
IM
NCM
p = 0.0293
p = 0.0225
I
Myeloperoxidase (MPO) staining: Colon: 11M ART
AL
CR (RM05)
100µm
100µm
Total MPO+ cells/mm$^2$
J
%HLA-DR+ CD4+ $T_m$
Blood
LN
K
%CD69+ CD4+ $T_m$
Blood
LN
L
Colon: 11M ART
%CXCR3+
CD4
p = 0.0075
M
%CXCR5+ PD-1+ $T_{fh}$
Colon
p = 0.0160
N
%CD25+ FoxP3+ $T_{reg}$
Colon
p = 0.1250

**Supplementary Figure 8. Caloric restriction (CR) promotes the sequestration of monocytes and $CD8^+$ T cells within the bone marrow.**

**(A)** Longitudinal graph depicting plasma viral load ($log_{10}$ SIV copies/mL) from infection to study conclusion. The shaded region marks the initiation of ART at 28 dpi. LOD = limit of detection.

**(B)** Viral decay rate (λ) quantifying the kinetics of plasma VL suppression below LOD following ART initiation.

**(C & D)** Expression of Bcl-2 (represented as MFI) in different $CD4^+$ **(C)** and $CD8^+$ **(D)** maturation subsets in the blood at 11M ART.

**(E & F)** Percentage of Caspase-3 expression in different $CD4^+$ **(E)** and $CD8^+$ **(F)** maturation subsets in the blood at 11M ART.

**(G)** Percentages of total $CD4^+$ and $CD8^+$ T cells in the bone marrow at 11M ART.

**(H)** Percentages of different monocyte subsets in the bone marrow at 11M ART.

**(I)** Representative image of IF staining for myeloperoxidase (MPO) in colon sections from AL and CR animals (left) and quantification of total MPO+ cells/$mm^2$ (right) at 11M ART.

**(J & K)** Percentage of $CD4^+$ T total memory T cells expressing HLA-DR **(J)**, and CD69 **(K)** in the blood and lymph nodes at 11M ART.

**(L)** Percentage of $CD4^+$ T cells expressing CXCR3 in the colon at 11M ART.

**(M)** %$CXCR5^+$ PD-$1^+$ $CD4^+$ $T_{fh}$ in the colon at 11M ART.

**(N)** %$FoxP3^+$ $CD4^+$ $T_{reg}$ cells in the colon at 11M ART.

**Statistical analyses: (B-N)** Mann-Whitney U test. α=0.05.

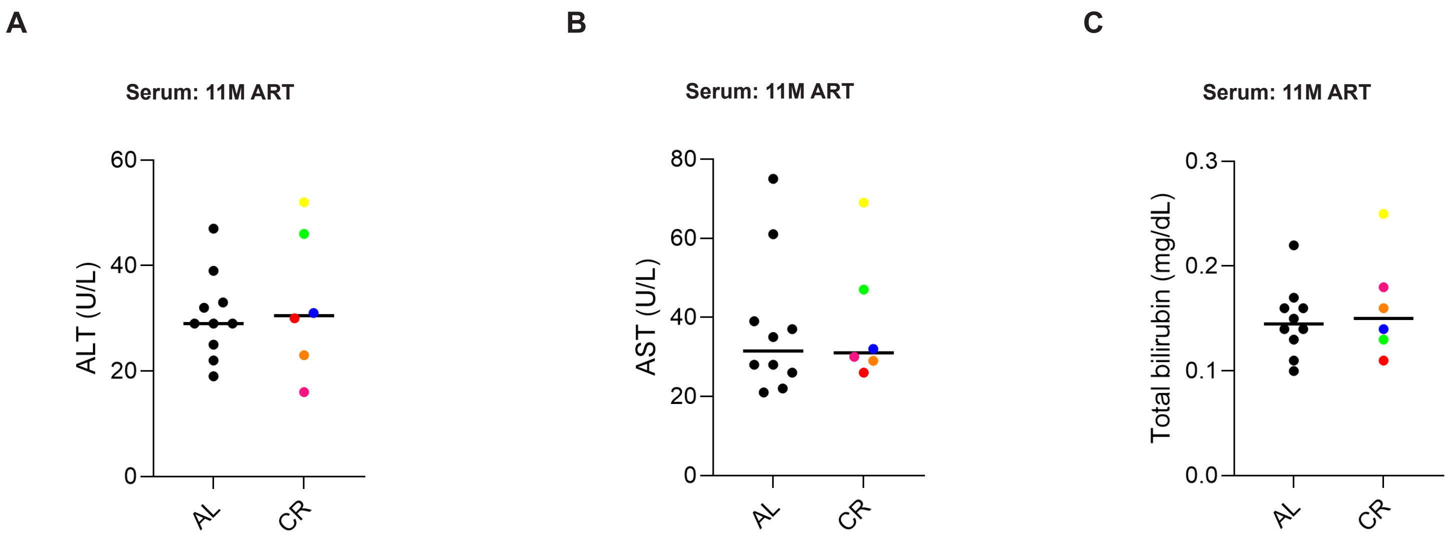


**Supplementary Figure 9. Comparable liver function in AL and CR animals following long-term ART.**

**(A-C)** Serum liver function tests in AL and CR animals measuring **(A)** ALT (alanine aminotransferase; U/L), **(B)** AST (aspartate aminotransferase; U/L), and **(C)** total bilirubin (mg/dL) at 11M ART.

**Statistical analyses: (A-C)** Mann-Whitney U test. α=0.05.

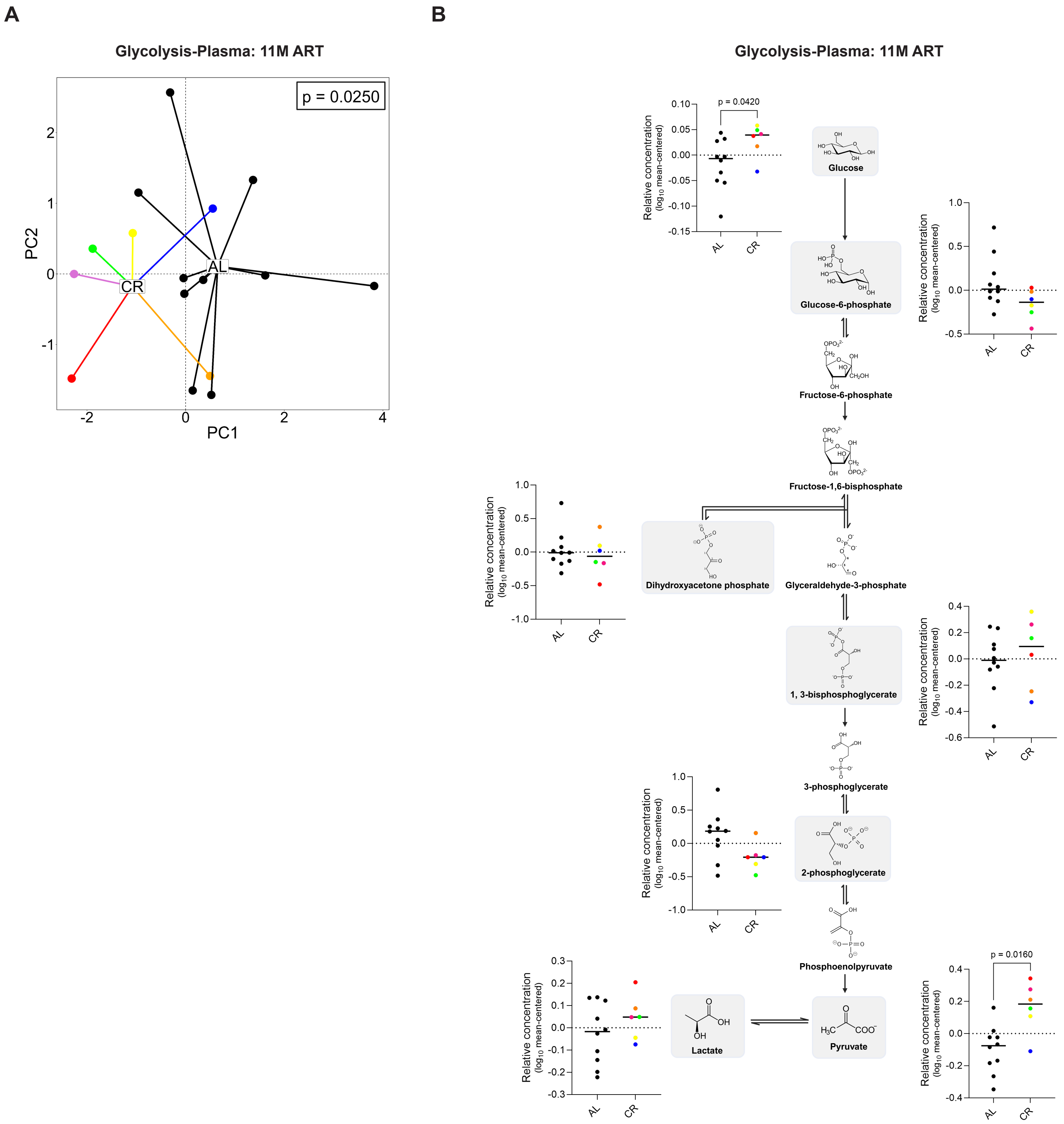


**Supplementary Figure 10. Metabolic reprogramming during ART results in minimal changes to the plasma glycolytic profile.**

**(A & B)** Principal component analysis (PCA) of key glycolytic metabolites **(A)** and their relative concentrations (mean-centered $log_{10}$ normalized) **(B)** in the plasma at 11M ART.

**Statistical analyses: (A):** PERMANOVA and **(B):** Mann-Whitney U test. α=0.05.

**Table S1: Median weight, body condition score, and levels of plasma physiological markers of animals after 4 months of CR versus the AL cohort prior to SIV infection.**

| Markers | AL | CR (30% CR) | p-value |
|---|---|---|---|
| **Weight (in kg.)** | 11.15 | 10.18 | 0.2250 |
| **Body Condition Scores** | 4 | 2.5 | 0.0061 |
| **Glucose (mg/dL)** | 56 | 65 | 0.0820 |
| **Cholesterol (mg/dL)** | 134 | 139 | 0.4043 |
| **Triglycerides (mg/dL)** | 54 | 54.5 | 0.5095 |
| **AST:ALT** | 1.31 | 1.01 | 0.1215 |
| **Hemoglobin (g/dL)** | 12.80 | 13.05 | 0.5732 |

**Statistical analyses:** Mann-Whitney U-test. $\alpha=0.05$.
